# Conformational free-energy landscapes of a Na^+^/Ca^2+^ exchanger explain its alternating-access mechanism and functional specificity

**DOI:** 10.1101/2023.01.20.524959

**Authors:** Fabrizio Marinelli, José D. Faraldo-Gómez

**Affiliations:** Theoretical Molecular Biophysics Laboratory National Heart, Lung and Blood Institute National Institutes of Health, Bethesda, MD 20814

## Abstract

Secondary-active transporters catalyze the movement of myriad substances across all cellular membranes, typically against opposing concentration gradients, and without consuming any ATP. To do so, these proteins employ an intriguing structural mechanism evolved to be activated only upon recognition or release of the transported species. We examine this self-regulated mechanism using a homolog of the cardiac Na^+^/Ca^2+^ exchanger as a model system. Using advanced computer simulations, we map out the complete functional cycle of this transporter, including unknown conformations that we validate against existing experimental data. Calculated free-energy landscapes reveal why this transporter functions as an antiporter rather than a symporter, why it specifically exchanges Na^+^ and Ca^2+^, and why the stoichiometry of this exchange is exactly 3:1. We also rationalize why the protein does not exchange H^+^ for either Ca^2+^ or Na^+^, despite being able to bind H^+^ and its high similarity with H^+^/Ca^2+^ exchangers. Interestingly, the nature of this transporter is not explained by its primary structural states, known as inward- and outward-open conformations; instead, the defining factor is the feasibility of conformational intermediates between those states, wherein access pathways leading to the substrate binding sites become simultaneously occluded from both sides of the membrane. This analysis offers a physically-coherent, broadly transferable route to understand the emergence of function from structure among secondary-active membrane transporters.

**Significance:** The class of membrane proteins known as secondary-active transporters mediate a wide range of critical cellular processes, including nutrient uptake, transmembrane signaling, and resistance to cytotoxic compounds, like human-made drugs. A detailed understanding of their molecular mechanisms is therefore of interest not only from a fundamental standpoint, but also because it will facilitate the design of inhibitors or stimulators that may be used as therapeutic agents. This study provides a conceptual mechanistic framework, grounded on statistical thermodynamics, that bridges the specific physiological function of these proteins and their molecular structure. While the study is focused on a particular subclass of transporters involved in cardiac physiology and cellular Ca^2+^ homeostasis, we envisage our conclusions will be broadly applicable.

## Introduction

Compared with other classes of membrane proteins, secondary-active transporters are arguably the least understood from a mechanistic standpoint, despite their pervasive role in all aspects of human physiology, both in health and disease. Like passive-diffusion channels and ATP-driven transporters, these proteins mediate the uptake or efflux of substances across membranes, in a selective manner, and to do so they cycle through a series of distinct structural states. However, unlike channels and primary-active transporters, this conformational cycle is neither initiated nor terminated by extrinsic factors, but rather by the recognition or release of the transported substances themselves, which can be thus thought of as both substrates and agonists. Another distinct characteristic of this class of proteins is their inherent poly-specificity, i.e. their structures have evolved the ability to selectively recognize substrates of different types, either concurrently (symporters) or competitively (antiporters). This feature is key, as it enables these proteins to harness a pre-existing transmembrane electrochemical gradient of one substrate, typically H^+^ or Na^+^ ions, to energize the accumulation or depletion of another, to a degree that would be unfeasible otherwise. Accordingly, cells employ this class of transporters for uptake of scarce nutrients and efflux of cytotoxic substances that penetrate the cell membrane, among other critical processes.

While much remains to be clarified about the interplay between secondary-active transporters and their substrates, the basic features of their conformational mechanism appear to have been delineated. Extensive biochemical and structural data demonstrate that these proteins undergo a spontaneous reversible transition between two major states, typically referred to as inward- and outward-facing (IF and OF) conformations; as a result, substrate binding sites in the protein interior become sequentially exposed to one or the other side of the membrane, but not both simultaneously (1–4). Meta-analyses of this class of transporters indicate this inherent conformational bi-stability owes to repeated topological units within their architectures, often inverted relative to the membrane plane (5); differences in the internal structure of these repeats naturally translate into asymmetric conformations for the transporter as a whole.

Despite these insights, fundamental questions about the nature of this ‘alternating-access’ mechanism remain to be answered. For example, the factors that control the probability of the interconversion between OF and IF states differ from protein to protein and are not at all self-evident from analysis of structures alone. For symporters, this interconversion occurs only when all substrates are simultaneously bound to the protein, or when all binding sites are simultaneously empty. Antiporters, by contrast, do not undergo this transition when no substrates are bound; instead, antiporters require that a substrate be recognized, but only of one type at a time. Furthermore, for both symporters and antiporters it is increasingly apparent that the transported substrate is not necessarily the substance that binds with the most specificity or potency; indeed, known inhibitors very often occupy the same binding sites as the biological substrates, and yet they somehow cause the alternating-access mechanism to stall.

Needless to say, the interconversion between OF and IF states is not abrupt; it entails a series of necessary intermediate conformations, whose nature and significance are also very much unclear. These intermediates have been rarely captured by conventional structural biology methods, indicating they are less energetically favorable than the OF and IF states, probably because they require the abovementioned structural repeats to adopt a similar arrangement, i.e. to form a quasi-symmetric conformation. At any rate, a logical expectation is that these intermediates will completely occlude access to the substrate binding sites in the protein interior, from either side of the membrane. The opposite scenario, i.e. simultaneous access to these sites from both sides, seems highly improbable as it would cause the dissipation of the electrochemical gradients that power active transport, as in a passive-diffusion membrane channel. An intriguing possibility, therefore, is that the mechanism by which substrates or inhibitors control the alternating-access transition somehow involves these occluded transient states that to date have been difficult to examine experimentally.

In this work we sought to obtain answers to these outstanding mechanistic questions, using as a model system a prokaryotic antiporter of Na^+^ and Ca^2+^, referred to as NCX_Mj (6–9). This transporter is a close homolog of NCX1, the human Na^+^/Ca^2+^ exchanger with a central role in the initiation and regulation of the heartbeat (10, 11). The primary function of NCX antiporters is to rapidly extrude cytosolic Ca^2+^, even when the concentration of free Ca^2+^ in the extracellular space exceeds that inside the cell. To minimize the backflow of Ca^2+^ into the cell in such conditions, this class of antiporters have co-evolved the ability to independently recognize and transport Na^+^ ions, which are more abundant in the extracellular space than in the cytosol. Thus, in typical physiological conditions the transporter captures cytosolic Ca^2+^ in the inward-facing state, delivers it extracellularly upon spontaneously transitioning to the outward-facing state, then loads Na^+^, and once again spontaneously transitions to the inward-facing state, where Na^+^ is released. (All these reactions are reversible, and some physiological conditions entail Ca^2+^ re-uptake and Na^+^ efflux instead.)

As is the case for many other transporters, however, the causality behind these observed processes is not understood at the molecular level. For example, it is unknown why the conversion between outward and inward-facing states requires that either Na^+^ or Ca^2+^ be bound; other monovalent and divalent ions bind to the transporter, such as H^+^, Li^+^, Cd^2+^ or Mn^2+^ and yet they are not transported, but rather inhibit function (6, 7, 9, 12, 13). It is also unclear why the alternating-access mechanism stalls when the protein is unliganded – which, as mentioned, differentiates antiporters from symporters. But perhaps the most intriguing question is why the stoichiometry of the antiport reaction mediated by both NCX1 and NCX_Mj is exactly 3 Na^+^ to 1 Ca^2+^ (9). That is, among the variety of partial or mixed ion occupancy states that necessarily exist, at least transiently, it appears that only those with either 3 Na^+^ or 1 Ca^+^ bound permit the transporter to spontaneously alternate between OF and IF conformations. It is worth noting that the exchange stoichiometry is a key physiological quantity, as it determines the thermodynamic limit for uphill Ca^2+^ efflux, for a given Na^+^ electrochemical gradient; for example, a 10-fold Na^+^ concentration gradient (larger outside) would allow NCX to deplete the cytosolic concentration of Ca^2+^ down to a value 1000 times smaller than that in the extracellular space. In other words, the exchange stoichiometry determines the intensity of the Ca^2+^ signals transduced by the transporter. To elucidate the molecular basis for such an important functional characteristic thus seems particularly worthwhile.

To clarify these rather fundamental questions, we have carried out a systematic single-molecule study of the structure and dynamics of NCX_Mj using advanced all-atom molecular-dynamics simulations; the trajectory time accumulated in this study, which exceeds 100 μs, illustrates the magnitude of the computational effort invested. Specifically, based on the only known structure of NCX_Mj, namely that of the OF state, we used enhanced-sampling simulation methodologies to map out the conformational free-energy landscape of the protein in a multi-dimensional space, and to examine how that landscape is reshaped by different ion occupancies including the apo state. These calculations enabled us to identify the structure of the IF conformation, as well as those of the most probable intermediates in the alternating-access mechanism, none of which had been known. Crucially, the calculated landscapes clearly explain why only the biological substrates catalyze this mechanism, but only for a unique stoichiometry. In summary, this study provides a conceptual mechanistic framework that bridges the specific physiological function of a paradigmatic human antiporter with its molecular structure and dynamics; we anticipate that this conceptualization will be readily transferable to other secondary-active transporters.

## Results and Discussion

### Asymmetry in topological repeats explains why known structure of NCX_Mj is outward-facing

All available evidence indicates that the known structures of NCX_Mj, obtained through X-ray diffraction, capture the architecture of the core ion-transporting domain of the proteins in the Ca^2+^-cation antiporter superfamily (TCDB database entry 2.A.190). These transporters are ubiquitous among eukaryotic and prokaryotic organisms, functioning primarily as either Na^+^/Ca^2+^ or H^+^/Ca^2+^ exchangers, in some instances modulated by other ions. This core domain contains a total of 10 transmembrane spans, all helical (**Fig. 1**). Closer examination reveals this assembly consists of two intertwined units of 5 helices each, whose transmembrane topology is inverted. The first repeat, TM1 to TM5, is fused to the second, TM6 to TM10, by a long, seemingly flexible linker. The amino-acid sequence identity between these two repeats is relatively low, however; in NCX_Mj, for example, it is only 32% (**Fig. S1**). Not surprisingly, therefore, the two repeats tend not to adopt exactly the same structure. Specifically, while helices TM3-TM5 and TM8-TM10 are organized almost identically, the configurations of TM1-TM2 and TM6-TM7 relative to the rest of each repeat are very different (**Fig. 1**). This difference translates into a noticeably asymmetry in the structure of the complete assembly. This asymmetry is very consequential; it opens up a pathway from the exterior of the protein into the sites where Na^+^, Ca^2+^ and other ions bind, deep in the interior. In the known structures of NCX_Mj, this access pathway is on the extracellular side of the protein; hence these structures are believed to capture the outward-facing state.

**Figure 1.**
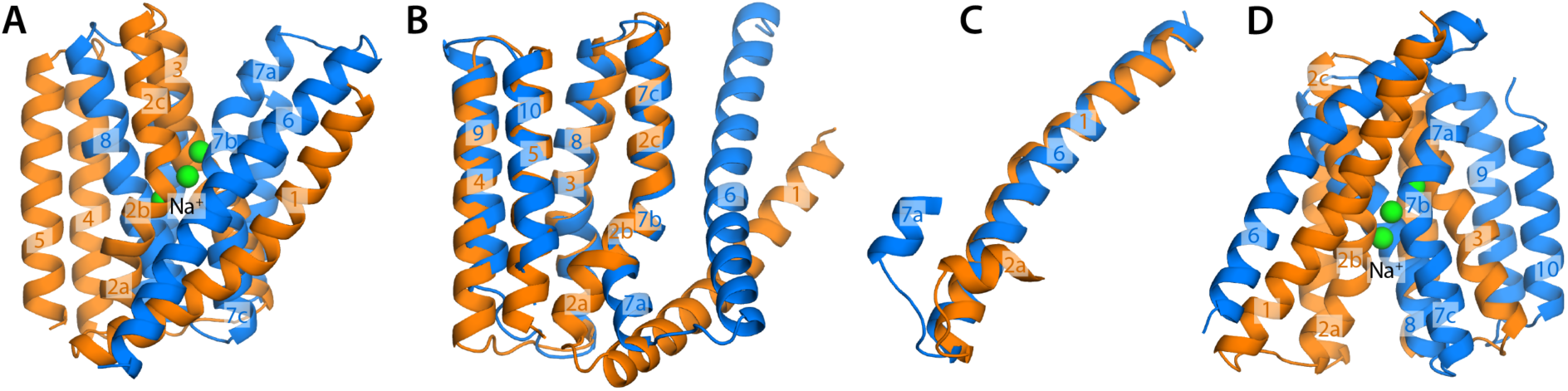
Topological repeats within the architecture of NCX_Mj, the Na^+^/Ca^2+^ exchanger from *M. jannaschii*. (**A**) Crystal structure of NCX_Mj in the outward-facing state (6). The structure contains two inverted topological repeats of 5 helices each, namely TM1-TM5 and TM6-TM10, shown in orange and marine, respectively. The protein is bound to three Na^+^ ions, shown in green. (**B-C**) Overlays of the internal structures of the two repeats reveals they are in part similar, but clearly distinct. The angle formed between TM2a and TM2b differs from that observed between TM7a and TM7b; the relative position of TM2a and TM1 also differs from that between TM7a and TM6. Taken together, these two differences explain why the structure shown in (A) features an access pathway into the Na^+^ binding site from the extracellular side. (**D**) A hypothetical structural model wherein the repeats swap conformations (see SI for more details), with no further adjustment, shows an intracellular access pathway analogous to that in panel (A).

An interesting exercise that further illustrates the mechanistic significance of this structural asymmetry is to generate a model of the transporter wherein the two repeats swap conformations (14). That is, in this repeat-swapped model, TM1-TM5 are arranged as observed for TM6-TM10 in the experimental structure of NCX_Mj, while TM6-TM10 are arranged as TM1-TM5 (see Supplementary Information for further details). Consistent with the logic outlined above, the resulting model features an access pathway into the ion binding sites on the intracellular side of the protein (**Fig. 1**). Needless to say, this simple model is unlikely to represent the actual structure of the inward-facing state, which to our knowledge has not been reported to date for this protein. However, it provides insights into the nature of the alternating-access mechanism, and a guide for a more rigorous characterization, as discussed in the following sections.

### Calculated free-energy landscapes reveal inward-facing structures bound to either Na^+^ or Ca^2+^

To identify, or at least predict, the actual structure of the IF state of NCX_Mj, we employed advanced molecular dynamics (MD) simulations. Specifically, we used an adaptive enhanced-sampling methodology known as a bias-exchange Metadynamics (15, 16). This approach is in our experience more capable than alternative MD simulation methodologies, for two important reasons: first, it permits a wide exploration of conformational space, even when this space is defined in multiple dimensions; and second, it permits a straightforward derivation of the free-energy landscape underlying that multidimensional space (as well as the error of the calculations) (8, 17, 18). In other words, as the simulation gradually reveals unknown conformational states, it is possible to use the principles of statistical thermodynamics to evaluate whether or not they are likely to be mechanistically significant. The simulations presented in this study used as the only input the X-ray structures of NCX_Mj bound to either 3 Na^+^ ions or 1 Ca^2+^ ion (6–8); however selected features of the abovementioned repeat-swapped model (**Fig. 1D**) guided some aspects of the calculation design (see Methods and **Fig. S2** for further details). As is common, a hydrated phospholipid bilayer was constructed around the protein to mimic in vitro experimental conditions (9). The simulations represented NCX_Mj, its ligands and environment in atomic detail (**Fig. S3A**). The trajectory time accumulated in these simulations was 62 μs.

The free-energy landscapes resulting from these calculations are shown in **Fig. 2A**. To facilitate the presentation of our results, the landscapes are projected onto a two-dimensional space, wherein one coordinate quantifies the degree of opening of the extracellular access pathway to the ion binding sites, while the other does the same for the intracellular pathway. For both Na^+^ and Ca^2+^, these maps reveal two distinct free-energy minima, i.e. two most probable states, separated by a less favorable region. Reassuringly, one of these minima encompasses the experimental structures of the outward-facing state of NCX_Mj bound to either 3 Na^+^ ions or 1 Ca^2+^ ion (6, 8). (Note no aspect of the calculation design dictates this result.) In this state, there are numerous contacts between helices TM1-2 and TM7-8 on the intracellular side of the protein, which close off the ion binding sites to the interior, while few contacts exist between TM2-3 and TM6-7 on the opposite side, which translate into an open exterior pathway (**Fig. 2B**). In the second free-energy minimum, these contact patterns are reversed almost exactly, and so the exterior pathway closes while an analogous interior pathway opens (**Fig. 2B**); that is, the calculated landscapes reveal the inward-facing structures of NCX_Mj loaded with either 3 Na^+^ or 1 Ca^2+^, which as mentioned had not been previously determined.

**Figure 2.**
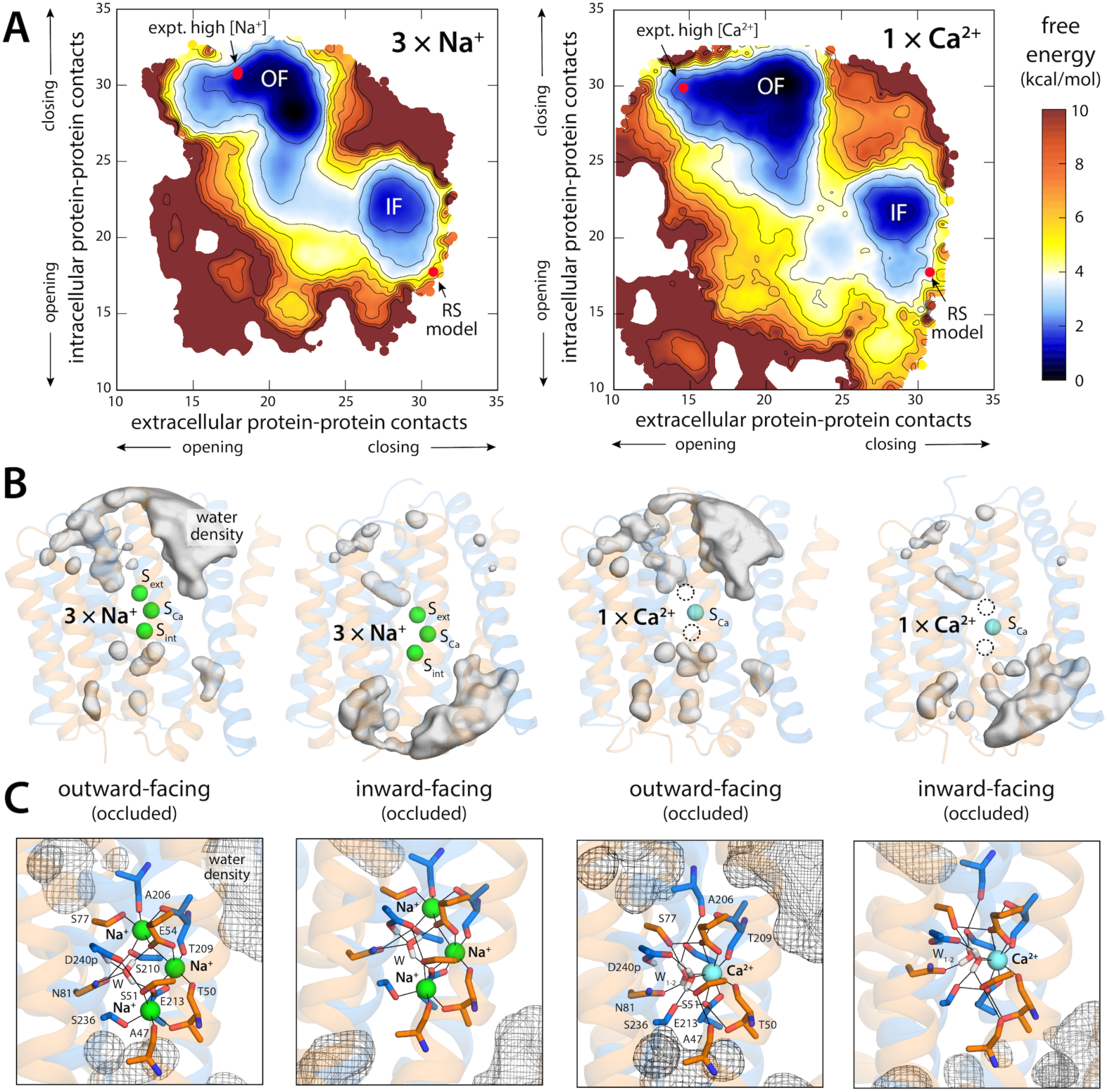
Free-energy landscapes underlying the alternating-access mechanism of NCX_Mj. **(A)** Landscapes are shown for the transporter loaded with 3 Na^+^ ions (top) or with 1 Ca^2+^ ion (bottom). For clarity, the maps represent the conformational free energy as a function of the degree of opening or closing of intra- and extracellular access pathways into the ion binding site. To objectively quantify accessibility, we use the number of contacts between two sets of protein residues lining those pathways. On the extracellular side, these contacts are between TM6 or TM7 and TM2 or TM3; on the intracellular side, between TM1 or TM2 and TM7 or TM8. Note that these selections are topologically symmetric with respect to the membrane plane. The two free-energy minima featured in each map correspond to the outward and inward-facing states (OF and IF) of the transporter. Red circles mark the positions in these maps of the outward-facing crystal structures of NCX_Mj obtained at high Na^+^ (left) or high Ca^2+^ (right) concentrations (PDB entries 3V5U/5HXE and 5HXR, respectively), and of the repeat-swap model shown in Fig. 1D. Contours are shown in intervals of 1 kcal/mol. Error estimates for each map are provided in **Fig. S3B**. (**B**) Water density maps for each of the free-energy minima revealed in the maps in panel (A) are overlaid onto representative configurations. For clarity only water molecules within 12 Å of the ion binding sites are mapped. Note OF and IF states have opposing water accessibility patterns, though in all cases the binding sites are occluded from the solvent. (**C**) Close-up of the ion binding sites, highlighting the mode of ion coordination in each case. Note the binding site geometries for OF and IF states are nearly identical.

Interestingly, the calculated free-energy maps clearly show that conformations wherein the binding sites are simultaneously exposed to both sides of the membrane (i.e. few or no protein-protein contacts on both sides of the structure) are energetically forbidden, consistent with the alternating-access model of active transport. Instead, the conversion between OF and IF entails a series of states wherein the number of protein-protein contacts closing off the intracellular and extracellular pathways is gradually reduced, but not to the extent required to make the binding sites accessible from either side (**Fig. 2A, 2B**). The nature of this conformational change will be discussed in more detail in the next sections. It is also worth noting that the detailed geometry of the ion binding sites is nearly identical when OF and IF states are compared, for both Na^+^ and Ca^2+^ (**Fig. 2C**). Consistent with these observations, conventional MD simulations initiated with configurations extracted from each of the free-energy minima revealed in the calculated landscapes showed minimal structural drift in the micro-second timescale; conversely, a conventional simulation of the repeat-swapped model shown above (**Fig. 1D**) showed a clear drift towards the minimum identified in the free-energy calculations (**Fig. S4**).

### Predicted IF state is consistent with HDX-MS data and recently-reported structure of NCX1

To validate or refute our prediction for the IF state of NCX_Mj, we first turned to a series of biochemical and biophysical experiments previously reported by Khananshvili and co-workers (19–21). These experiments compared two forms of NCX_Mj: a WT-like construct, which structural studies had shown to strongly favor the OF state when solubilized in detergent (6, 8); and a functional mutant known as 5L6-8 (due to an elongation of the TM5-TM6 loop), which transport assays had indicated also populates the IF state (19). Consistent with those studies, Khananshvili and co-workers observed that cysteines engineered in positions G42 and G201, in the intracellular and extracellular sides of the protein, respectively, showed opposing reactivity patterns to TMRM maleimide probes introduced on either side of the membrane (21). As shown in **Fig. 3A**, these observations are in line with our prediction; G42 is maximally exposed to the solvent in the predicted IF state while G201 is buried within the protein, while the opposite is true for the OF state. Interestingly, the conformational intermediates show a reduced but equivalent accessibility for both positions, as might be expected for quasi-symmetric states.

**Figure 3.**
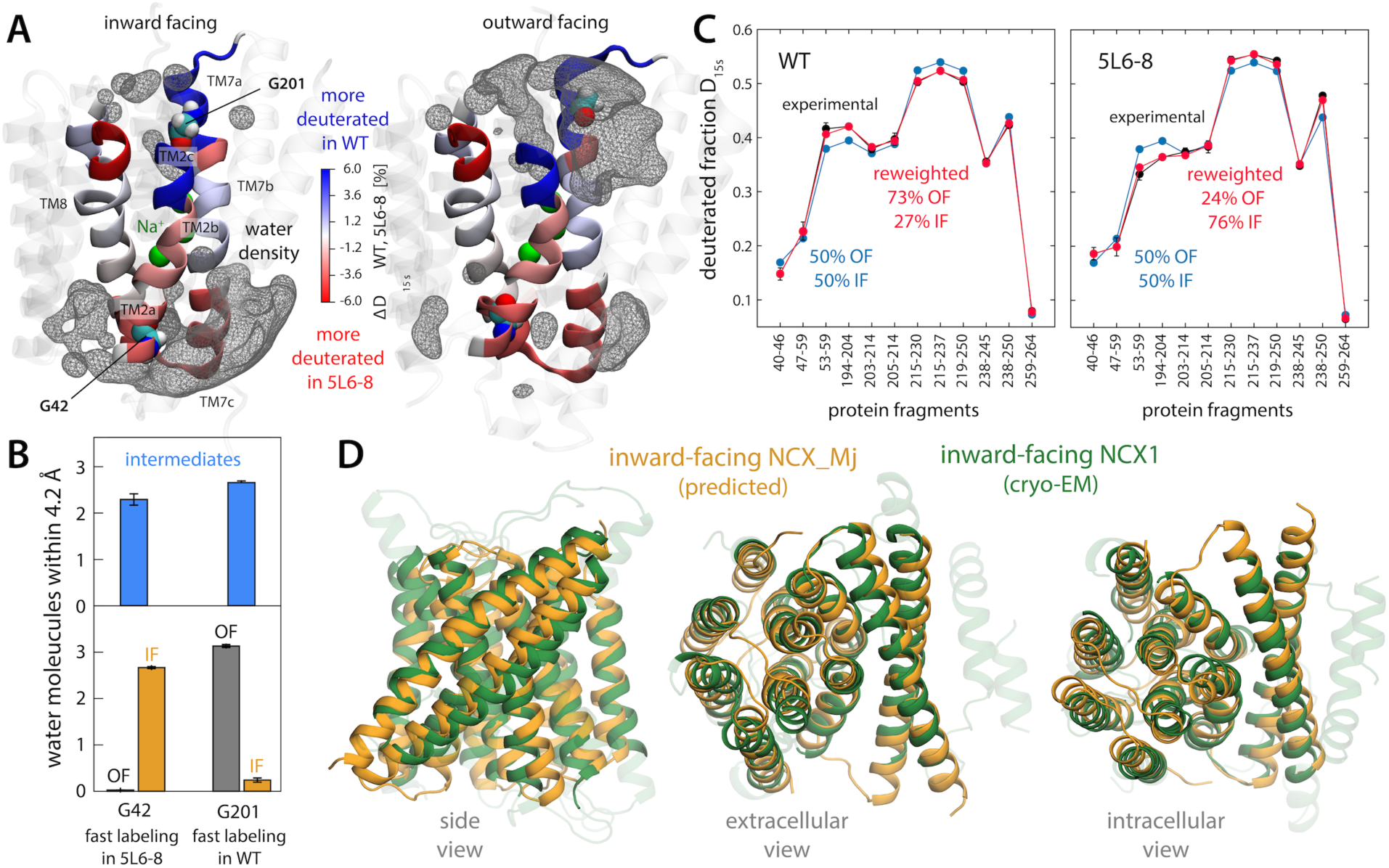
Validation of the predicted IF structure of NCX_Mj against available experimental data. (**A**) Graphical representation of published HDX-MS data (*21*) for two forms of NCX_Mj (WT and 5L6-8) with distinct propensities to adopt the OF of IF state (see text). Differences in measured deuteration levels for these two forms (after 15 s and for saturating [Na^+^] conditions) are indicated with a color scale, for the regions of the protein that were examined experimentally (TM2, TM7 and TM8). Data on overlapping protein fragments has been broken down into smaller regions by linear combination of the deuterated fractions. Residues G42 and G201, whose accessibility was probed through complementary labeling assays, are also highlighted. For reference, the figure also shows density isosurfaces for water molecules within 12 Å of the protein binding sites (gray mesh), based on our simulation data. (**B**) Solvent accessibility of G42 and G201 in either the OF or IF states identified in the free-energy maps in Fig. 2, as well as in the intermediate regions. This accessibility is quantified by the number of number water molecules found, on average, within 4.2 Å of either residue. (**C**) For both WT and 5L6-8, the degree of deuteration measured for a collection of protein fragments in TM2, TM7 and TM8 (after 15 s and for saturating [Na^+^]) is contrasted with deuteration levels calculated for an ensemble of OF and IF structures extracted from the free-energy basins in Fig. 2 (see Supplementary Information). Calculated and experimental data are compared when OF and IF are equally weighted, and for alternative weights that result in optimal agreement with measured data, for either WT or 5L6-8 (see also Fig. S5). These population shifts are consistent with the known conformational propensities of these forms. (**D**) Comparison between the IF state of NCX_Mj identified in the free-energy landscapes shown in Fig. 2 with a recently reported cryo-EM structure of human NCX1 (23), coincidentally captured in the IF state. The RMS difference between the Cα traces is 1.9 Å.

Khananshvili and co-workers also compared the degree of hydrogen-deuterium exchange for WT and 5L6-8 NCX_Mj in various conditions and for multiple D_2_O exposure times (20, 21). The rate of this exchange is a complementary metric of solvent exposure (among other factors). The data obtained for saturating Na^+^ concentrations is particularly relatable to our simulation conditions, in that the binding sites within the transporter ought to be fully (and solely) occupied by Na^+^; likewise, only exposure times shorter than 15 s seem pertinent, as beyond that point local protein unfolding likely becomes increasingly dominant. To evaluate whether or not our simulation results are consistent with this data, we employed a variation of an analysis methodology we recently reported, named HDXer, based on the maximum-entropy principle (22) (see Methods and Supplementary Information for further details). In a nutshell, this methodology considers an ensemble of protein conformations that includes two or more distinct populations (obtained through MD simulations or any other modeling method) and determines what population shifts would be required for that ensemble to optimally match an input HDX dataset. (Note that for this optimization to result in a good match or plausible population shifts, much of the underlying sample must be compatible with the experimental data.) Following this approach, we extracted an ensemble of OF and IF conformations from the free-energy minima identified in the calculations reported in **Fig. 2A**, populating each state equally. We then used the new version of HDXer to calculate the deuteration rates expected for the set of protein fragments examined experimentally, and to gradually reweight the initial populations of OF and IF states so as to obtain maximal agreement with experiment. As shown in **Fig. 3C** (and **Fig. S5**), we found that plausible population shifts resulted in excellent agreement with the experimental data. Specifically, in order to reproduce the HDX dataset for the WT protein, the population of the OF state has to be upshifted, in line with the fact this form of the protein favors this particular state in experimental conditions. By contrast, reproducing the HDX dataset for the 5L6-8 mutant requires a depletion of the OF state and a larger population of the IF state, which is again in line with the experimental characterization of this mutant. In summary, while recognizing that HDX measurements offer limited structural resolution, we believe this objective analysis of the experimental data lends confidence to our prediction of the IF state of NCX_Mj.

As this manuscript was in its final stages of preparation, Jiang and co-workers reported the first known structure of NCX1, the human homolog of NCX_Mj found in cardiac muscle (23). As for other membrane proteins, complexation with an antibody fragment appears to have been crucial for this breakthrough and coincidentally, this approach captured the transporter in what appears to be the IF state. As demonstrated in **Fig. 3D**, our prediction for the IF of NCX_Mj is highly consistent with this new structure. Notwithstanding the low sequence identity between these proteins (∼20%), it is evident from an overlay of the two structures that they represent the same functional state (RMS difference of 1.9 Å based on aligned Cα atoms). IF structures have also been reported for several H^+^/ Ca^2+^ exchangers (24–26). Those structures are also comparable though logically not identical to that predicted here for NCX_Mj; for example, for CAX_Af (∼20% sequence identity), the RMS difference between the structures is only 2.1 Å.

### Alternating-access transition entails a new type of structural mechanism

Having validated our prediction for the IF state of NCX_Mj, we proceed to analyze in more detail the structural and energetic features of the mechanism of alternating access in NCX_Mj. Overlay of representative OF and IF structures underscores the significance of the structural asymmetry between the two topological repeats within the protein (**Fig. 4A, 4B**); only the elements that distinguish these repeats in the experimental structure of the OF state, i.e. TM1, TM2, TM6, TM7, change upon transition to the IF state, while the rest of the protein remains largely unchanged. The resulting mechanism is one wherein TM1-TM6 travels across the membrane midplane by about 1 nm, accompanied by more subtle changes in the intracellular and extracellular portions of TM2 and TM7; by contrast, the geometry of the ion binding sites and their position relative to the membrane are unaltered (**Fig. 4A, 4B**). To our knowledge, this mechanism is unlike those previously described for other active transporters, though there are some commonalities with the so-called rocker-switch model (1–4).

**Figure 4.**
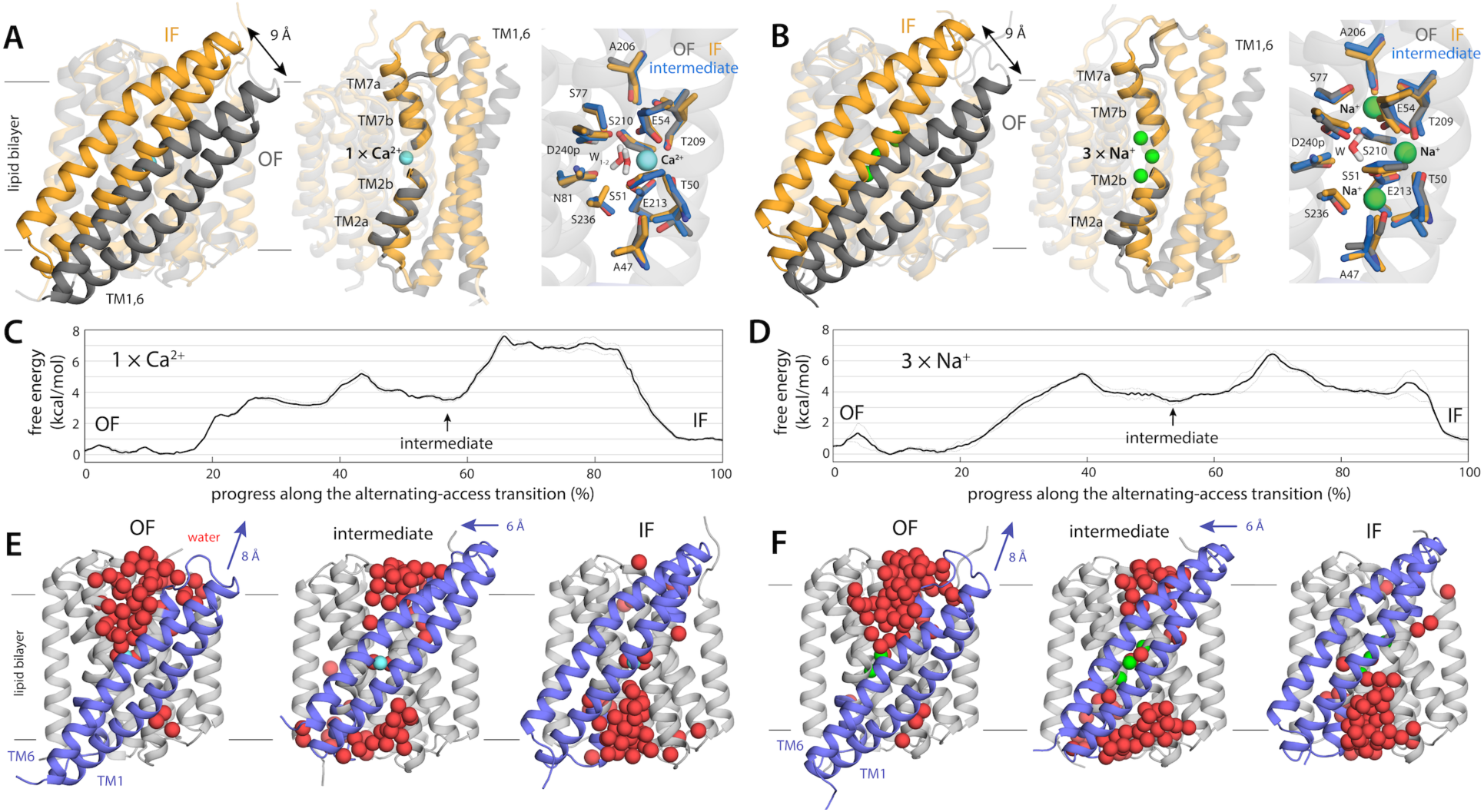
Mechanism of alternating-access in NCX_Mj inferred from analysis of simulation data. (**A, B**) Comparison between the OF and IF states, using representative configurations of the free-energy minima in Fig. 2, for the Ca^2+^- and Na^+^-bound transporter, respectively. Note the displacement of the TM1-TM6 unit across the membrane midplane, alongside localized changes in the intracellular and extracellular halves of TM2 (TM2b) and TM7 (TM7ab), respectively. Close-ups of the ion binding sites are also shown, for the OF and IF states as well as for a doubly-occluded intermediate (see below). (**C, D**) Change in free-energy along the minimum free-energy (most probable) multi-dimensional path connecting the OF and IF states, for the Ca^2+^- and Na^+^-bound transporter, respectively. The profiles reveal a metastable intermediate about halfway through the transition. (**E, F**) OF, intermediate and IF states are compared side by side, highlighting water molecules in proximity to the ion-binding sites, filling access pathways into the protein interior from either side of the membrane. Note the displacement of TM1-TM6 in panels (A, B) in fact entails two distinct movements in different directions, as indicated.

To adequately characterize this mechanism, however, we must go beyond this comparison and examine the intermediates of the transition. To do so, we re-analyzed our simulation data to identify the minimum free-energy pathway connecting the OF and IF states, i.e. the most probable trajectory followed by the transporter as it undergoes this conformational change (note this pathway is multi-dimensional and thus not discernible in the 2D projection used in Fig. 2; see Methods for further details). Key features of this structural mechanism are shown in **Fig. 4C-4F** (see also **Movies S1-S2**, and **Fig. S6-S8**). Starting from the OF state, the first stage of the transition entails displacement of TM1-TM6 toward the extracellular space and a change of TM2a from a sustained π-helical configuration to a more labile state (including α-helical, π-helical and partially unstructured conformations); the change in TM2a requires disruption of a hydrogen-bond between G42 and A47 and displacement of F39 from a protected hydrophobic pocket. These changes permit water to begin to penetrate the structure from the intracellular side, while the access pathway on extracellular side begins to be depleted. However, water exposure of hydrophobic intracellular residues and disruption of several salt-bridges on that side of the protein (R223-E28, R223-D21) lead to a sharp increase in free energy, peaking at around 5 kcal/mol. Soon thereafter, however, the protein settles down into a metastable state through a reconfiguration of sidechains and the surrounding solvent; at this point, TM1 and TM6 have moved toward the extracellular side by about ∼8 Å, relative to the OF state (**Fig. 4E, 4F**), and the ion binding sites are simultaneously occluded from both sides of the membrane. The next step entails crossing the highest free-energy barrier of the transition, which is ∼7.5 kcal/mol when Ca^+^ is bound state and ∼6.5 kcal/mol when 3 Na^+^ ions are carried instead. In this step, TM1 and TM6 continue to move, but this motion is now parallel to the membrane plane and requires reconfiguration of protein-protein and protein-lipid interactions involving bulky hydrophobic residues on both sides of the protein (e.g. F23 and F182). A second intermediate forms thereafter, and at this point TM1 and TM6 are essentially in their final position. However, complete progression towards the IF state requires a rearrangement of TM7a that mirrors the changes seen earlier in TM2a, but in reverse; i.e. TM7a becomes π-helical, as a hydrogen-bond between G201-A206 form and F202 becomes enclosed in a hydrophobic pocket. A substantial reduction in free-energy is then achieved through dehydration of the extracellular access pathway and the formation of several salt-bridges (K198-E257, K187-D197) on that side of the protein.

It is apparent from this analysis that the conformational change underlying the alternating mechanism of NCX_Mj would be very poorly described by a linear geometric interpolation between the OF and IF states. Instead, this mechanism entails a complex choreography of spontaneous progressive rearrangements, both in the protein and its local environment, which ultimately translate into larger-scale changes in tertiary structure. Interestingly, as the protein changes, we observe no significant effects on membrane shape or thickness (**Fig. S9**), despite the motion of TM1 and TM6 across the lipid-bilayer midplane and the opening and closing of water channels near lipid-solvent interface; this observation is in line with the fact that the turnover rate of NCX_Mj is rather insensitive to the lipid composition of the membrane (27). In this regard, therefore, NCX_Mj is unlike other transporters that deform the membrane as a result of large domain motions perpendicularly to the bilayer plane (28), probably because the TM1-TM6 unit is comparatively much smaller.

As mentioned, the binding site region also retains the same structure throughout the transition (**Fig. 4A, 4B**); we can thus infer that protein-substrate interactions, once fully engaged, are no longer a determining factor in the energetics of the transition between the OF and IF states. What other interactions might be more impactful? To examine this question, we recalculated the free-energy differences between the OF state and either the IF or the intermediate states, by resampling and reweighting our simulation data with modified versions of the original forcefield wherein the atomic charges of a selected number of residues are slightly scaled down (**Fig. S8**). The rationale behind this analysis is that this modification of the energy function impacts both protein-protein and protein-solvent interactions, thus informing on the global electrostatic contribution of the selected residues. First, we considered a selection that includes all the polar residues near the ion-binding sites (those highlighted in **Fig. 2C**) as well as the ions themselves. Consistent with the inference made above, we found that scaling down the magnitude of these interactions had no significant effect on the free-energy difference between the OF and IF state, or that between the OF and intermediate states, relative to the original calculation with an intact forcefield. By contrast, perturbation of more peripheral polar interactions had a pronounced effect; relative to the OF state, the free-energy of the intermediate is significantly diminished when salt-bridges and other residues engaged in protein-protein or protein-water hydrogen-bonds are weakened. Interestingly the free-energy difference between the OF and IF states are also affected. These results underscore the mechanistic significance of these peripheral regions, often overlooked because they are less conserved.

Indeed, it is worth noting that, as is often observed for other transport families, NCX_Mj is markedly slower than its eukaryotic homologs, like cardiac NCX1 (29–31). The reasons are unclear, but we believe that the methodology deployed in this study, when applied to NCX1, will reveal the origins of this important difference. Distinct protein-protein and protein-solvent interaction patterns in peripheral regions of the protein are in our view the most likely explanation, rather than distinct protein-substrate interactions.

### Functional specificity is also encoded in the free-energy landscape of the transporter

Three characteristics summarize the functional specificity of a secondary-active transporter. First, what is its mode of transport? That is, does the protein function as an antiporter, a symporter or a uniporter? Second, what are substrates that are translocated across the membrane, among all possible substances that might transiently bind to the protein? And third, what is the precise stoichiometry of the transport cycle? To anticipate or rationalize the answer to these questions on the basis of molecular structure is one of the central, largely unresolved problems in this field of research. For many transporters, these functional characteristics had been established through biochemical assays well before their molecular structure became known, and so the nature of this structure-function relationship, when examined post-hoc, might appear straightforward. Even then, however, it can be argued that to convincingly deduce this relationship is not at all trivial; this difficulty is illustrated by many other cases wherein a transporter structure has been determined, typically that of an obscure prokaryotic homolog, before comprehensive biochemical assays exist. In those cases, to anticipate the functional specificity of the transporter is clearly not possible solely based on inspection of molecular structures. We reasoned, however, that analysis of conformational landscapes calculated for different occupancy states might provide answers to the questions outline above, in the case of NCX_Mj.

Following that logic, we first tried to understand why NCX_Mj does not function as a Na^+^ or Ca^2+^ uniporter (or symporter), i.e. why translocation of one of these types of ions is necessarily coupled to the other. A uniporter mode, however, would require that the transporter undergo the alternating-access transition when neither Na^+^ or Ca^2+^ are bound (similarly for a symporter). To examine whether that is the case, we removed these ions from the OF and IF structures revealed by our free-energy calculations, and recalculated the free-energy landscape of the transporter in this apo state, using the same methodology used previously. As shown in **Fig. 5A**, this new landscape is radically different from that observed in the presence of 3 Na^+^ or 1 Ca^2+^. While the landscape still features an OF and an IF state, it is effectively impossible for the transporter to interconvert between those two states. Furthermore, the OF and IF states are distinctly unlike those observed when 3 Na^+^ or 1 Ca^2+^ are bound. In the absence of these ions, the binding site region is widely open to either the extracellular or intracellular solution (**Fig. 5B, 5C**); indeed, comparison of the maps shown in **Fig. 2** and **Fig. 5** makes it clear that the occluded OF and IF states that are favored when 3 Na^+^ or 1 Ca^2+^ occupy the protein, and which precede the alternating-access transition, are energetically unattainable in this apo state, seemingly because the set of charged and polar sidechains that would be otherwise involved in ion coordination (e.g. E54 and E213) cannot reconfigure so as to eliminate unpaired groups, making dehydration of this region inviable. Hence, that NCX_Mj does not function as a uniporter ultimately owes to the unfeasibility to this occlusion process.

**Figure 5.**
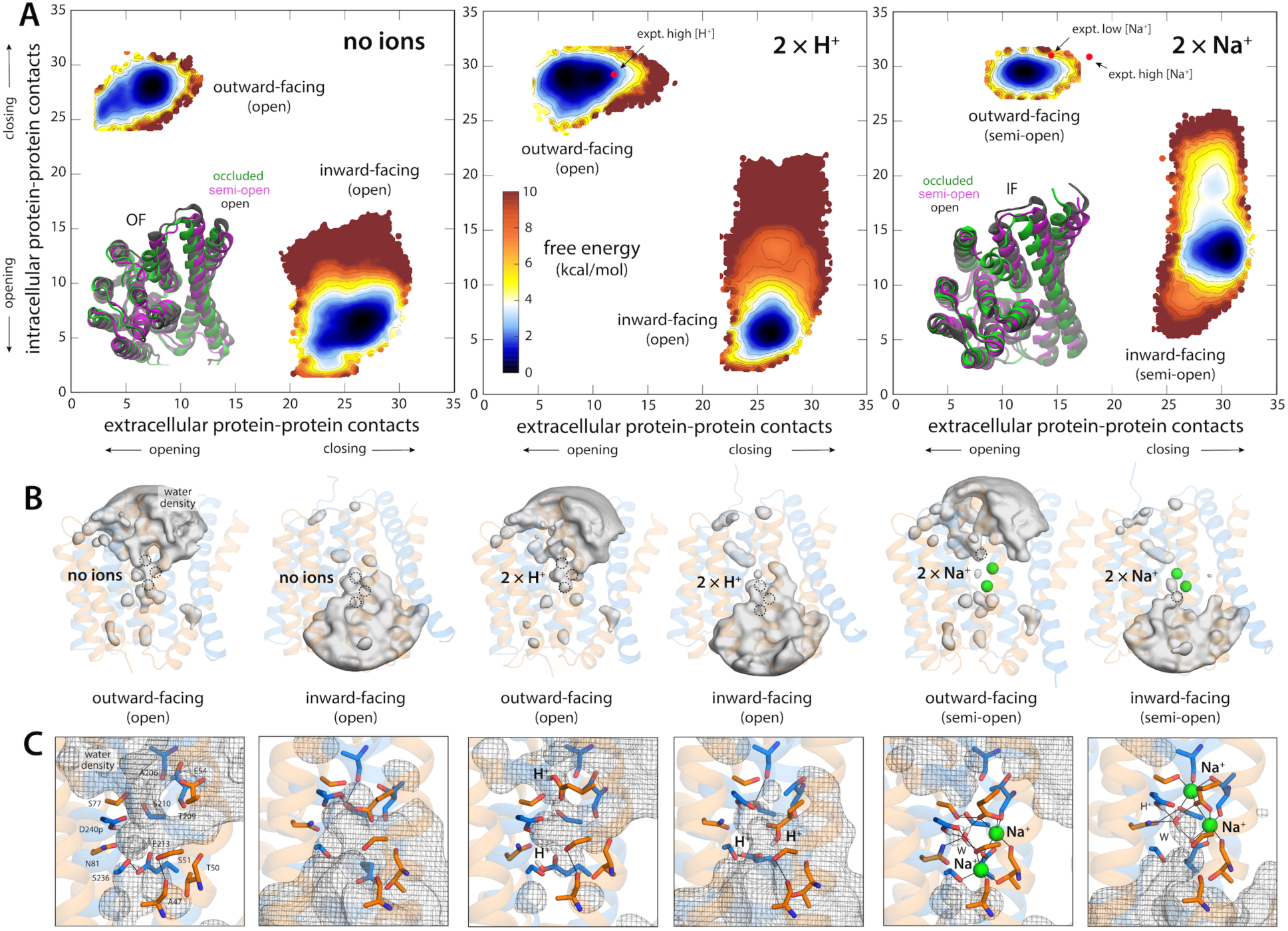
Free-energy landscapes dictate the functional specificity of NCX_Mj. (**A**) Free-energy landscapes analogous to those shown in Fig. 2, but for different ion-occupancy states, namely with no ions bound, with 2 H^+^ bound to E54 and E213, and with 2 Na^+^ bound only. Red circles mark the positions in these maps of the outward-facing crystal structures of NCX_Mj obtained at low pH and at low and high [Na^+^] (PDB entries 5HXH, 5HWY and 5HXE, respectively). The landscapes show the alternating-access transition is energetically unfeasible, and that the transporters are trapped either in OF or IF conformations that are distinctly more open to the solvent that those favored when either 3 Na^+^ or 1 Ca^2+^ ion are bound. Insets show an overlay of these different conformations, for both the OF (left) and IF (right) states. Error estimates for each map are provided in **Fig. S3C**. (**B**) Water density maps for each of the free-energy minima revealed in the maps in panel (A) are overlaid onto representative configurations. For clarity only water molecules within 12 Å of the ion binding sites are mapped. Note OF and IF states have opposing water accessibility patterns, and that in all cases the ion binding sites are readily exposed to the solvent, but only on one side of the membrane. (**C**) Close-up of the ion binding sites, highlighting the configuration of the ion coordination shell in each case.

As mentioned, the family of calcium/cation exchangers that includes all NCX proteins also includes a wide array of H^+^/Ca^2+^ exchangers, whose structures are highly similar (in the core transport domain). NCX_Mj, however, does not exchange Ca^2+^ for H^+^, nor does it exchange Na^+^ for H^+^ either (9). The reason is not that H^+^ do not bind to NCX_Mj; indeed, in conditions where the concentration of H^+^ is comparable to that of Na^+^ or Ca^2+^ (i.e. low pH), NCX_Mj binds H^+^, through protonation of E54 and E213, and becomes inhibited (7). A free-energy landscape calculated for this condition explains why this is case. As noted for the apo state, the landscape demonstrates that occlusion of the binding sites in either the OF or the IF states is effectively unfeasible, and by extension, so is the alternating-access transition (**Fig. 5A**). That occlusion is so energetically unfavored despite the neutralization of E54 and E213 underscores that dehydration of the binding site region is unlikely without a well-defined interaction network wherein all polar groups are paired (**Fig. 5B, 5C**); the net charge of the binding site is clearly of little or no consequence, and is thus not a good predictor of the viability of the alternating-access transition.

Lastly, we examined why the ion-exchange stoichiometry of NCX_Mj (and by extension its close homologs like cardiac NCX1) is precisely 3 Na^+^ for 1 Ca^2+^ (9), which as mentioned dictates the maximum concentration gradient against which the transporter will continue to drive Ca^2+^ efflux, for a given inward Na^+^ gradient (also known as sodium-motive-force). In particular, we examined why the nearest possibility, namely 2 Na^+^ for 1 Ca^2+^, is not viable (9). To do so, we removed one of the 3 Na^+^ ions from the OF and IF occluded structures (keeping in each case the 2 ions that are deeper into the binding site), and recalculated the free-energy landscape of the transporter. Like for the apo transporter and the H^+^-bound state, the landscape features clearly defined OF and IF states, but also demonstrates the interconversion between these states is effectively not possible (**Fig. 5A**). Interestingly, though, with 2 Na^+^ ions bound the binding site favors a configuration that is partially occluded, i.e. while the empty binding site remains exposed to the solvent, the self-organization of the other two sites occupied by Na^+^ appears to greatly facilitate their dehydration (**Fig. 5B, 5C**). Nevertheless, it seems clear from our data that full occlusion, and therefore the initiation of the alternating-access transition, requires recognition of the third Na^+^ ion.

In summary, this analysis demonstrates that the functional specificity of a transporter can be in fact predicted or rationalized from structure alone, with limited a priori input from biochemical data. It is however important to recognize that purely visual or heuristic analyses of individual structures are insufficient to bridge the gap between structure and function, as are cursory simulation studies based on anecdotal data. By contrast, we posit that conformational free-energy landscapes, calculated with carefully-designed advanced simulation methods, are a highly promising route.

## Conclusions

Overwhelming evidence indicates that the alternating-access model is the most plausible framework with which to rationalize secondary-active membrane transport at the molecular level (i.e. uphill transport without ATP hydrolysis). It is underappreciated, however, that this model rests on two seemingly paradoxical principles, which deserve further scrutiny. First, that the interconversion between OF and IF conformations, or vice versa, is not driven by extrinsic forces, such as transmembrane voltages, concentration gradients or binding-affinity differentials. Instead, this interconversion is spontaneous and stochastic, and perfectly reversible. Second, that despite this inherent plasticity, this interconversion occurs with measurable likelihood only for two concrete substrate-occupancy states among all those that are possible, irrespective of whether these states are transient or long-lasting. Indeed, the nature of these two specific occupancy states defines whether the protein functions as a symporter, an antiporter or a uniporter; it also defines how many and what type of substrates are actually translocated across the membrane, which in turns determines the maximum transport capacity of a given transporter population, and the energy source that sustains their biological activity.

In this study, we show that the theory of conformational free-energy landscapes provides a means to reconcile these key requirements. Using the Na^+^/Ca^2+^ exchanger NCX_Mj as a model system (**Fig. 6**), we show that the expected OF and IF conformations (the latter of which was previously unknown) are indeed the two most prominent states in the free-energy landscape of transporter, and show these states reflect two alternate configurations for each of two topological repeats within the protein architecture. The calculated landscapes also indicate that in many conditions the free-energy barrier between the OF and IF states will be effectively unsurmountable, at least in the physiological timescale. For example, when no ligands are bound to the transporter or when the transporter is loaded with H^+^, the transporter is arrested in either the OF or IF state, but will not interconvert between those states. By contrast, recognition of either 1 Ca^2+^ ion or 3 Na^+^ ions radically re-shapes the free-energy landscape of the protein; a series of conformational intermediates between the OF and IF states become energetically attainable, which facilitate a gradual interconversion between those states simply through thermal fluctuations of the molecular system.

**Figure 6.**
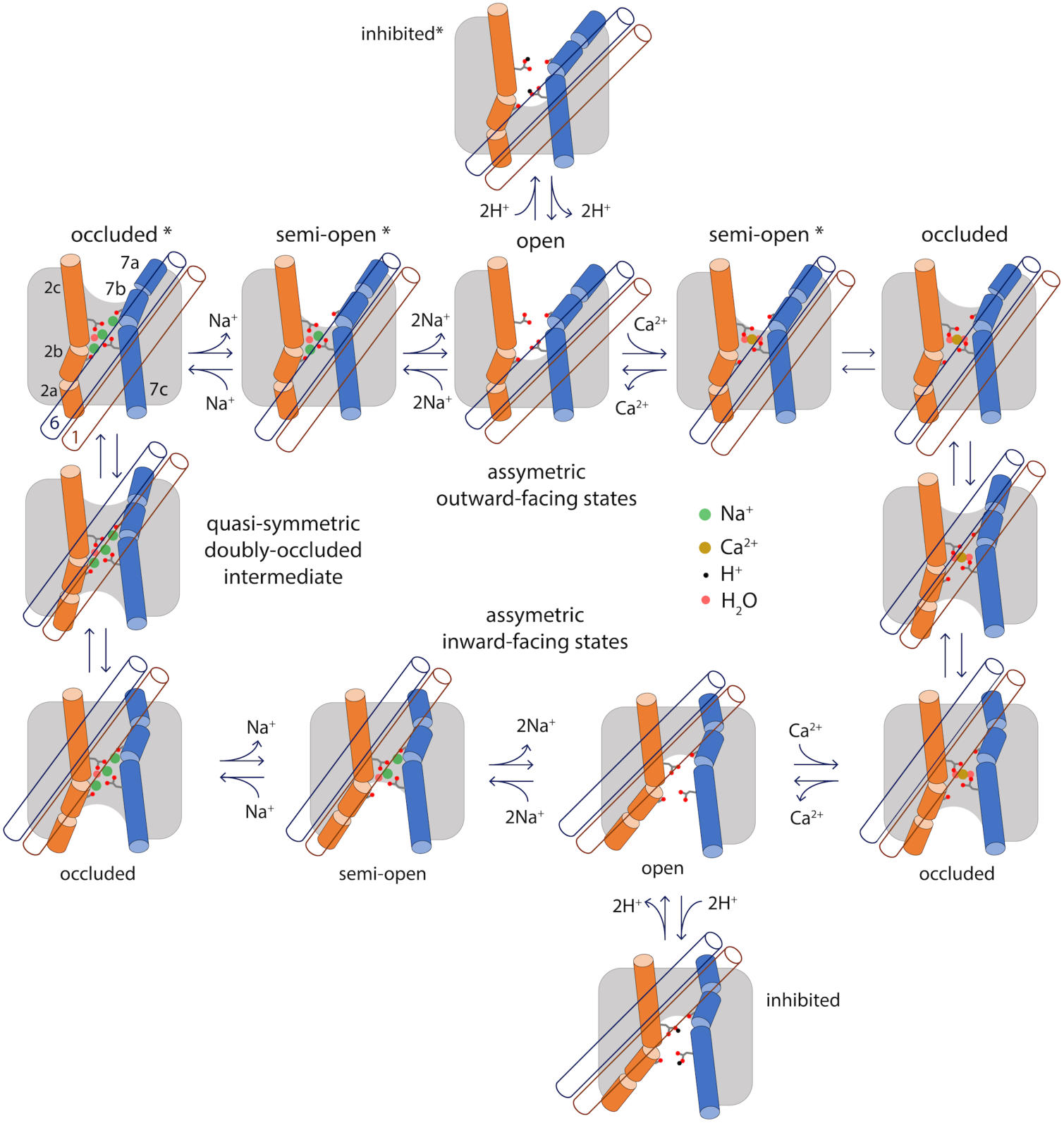
Specificity in the mechanism of alternating-access of the Na^+^/Ca^2+^ exchanger. This study demonstrates that the functionality of NCX_Mj, namely antiport of 3 Na^+^ and 1 Ca^+^, owes to the fact that the alternating-access transition is not viable for any other substrate-occupancy states, including the apo protein. The scheme highlights the essence of the conformational changes that the transporter undergoes when transitioning between the OF and IF states, namely a sliding motion of TM1 and TM6 (blue and brown empty cylinders, respectively) across the membrane mid-plane, and changes in secondary structure in TM2 and TM7 (orange and marine full cylinders, respectively). Also highlighted are the side chains of E54 (on TM2c) and E213 (on TM7c) and the carbonyl groups of A47 (on TM2b) and A206 (on TM7b), which coordinate Na^+^, Ca^2+^ and H^+^. The scheme shows only the states and connectivity deduced from the calculated free-energy landscapes in Fig. 2 and Fig. 5A; additional open or semi-open states with only 1 Na^+^ or 1 H^+^ bound, or a combination of Na^+^, Ca^2+^ and H^+^, are also conceivable. Asterisks mark the states that have been also determined by X-ray crystallography.

In this view, it becomes clear why the alternating-access mechanism is a spontaneous, reversible process that requires no extrinsic driving force, and yet one that is tightly controlled. In a hypothetical condition where free Ca^2+^ is similarly available across the membrane, but Na^+^ is much more abundant extracellularly, a population of NCX_Mj transporters will gradually deplete the cytosol from Ca^2+^, i.e. generate an outward electrochemical gradient of Ca^2+^ that ultimately balances that of Na^+^. However, this Na^+^ gradient does not cause or drive the protein towards one or other conformational state; it merely introduces a statistical bias or preferred directionality in the alternating-access cycle. Specifically, most (but not all) of the transitions from the OF to the IF state will occur with 3 Na^+^ ions bound, and many (but not all) of the transitions from the IF to the OF state will instead occur with 1 Ca^2+^ bound. Consistent with this notion, in the absence of Ca^2+^ NCX_Mj will not catalyze a net uptake of Na^+^, even under a strongly favoring electrochemical gradient; instead, the transporter will repeatedly interconvert between OF and IF states, but only with 3 Na^+^ ions bound, but not when empty, and therefore the Na^+^ concentrations at either side of the membrane will remain unchanged. Similarly, even in pH conditions such that H^+^ outcompete Na^+^, an inward gradient of H^+^ will not result in Ca^2+^ efflux despite the many structural similarities between Na^+^/Ca^2+^ and H^+^/Ca^2+^ exchangers, as the binding of H^+^ in fact precludes the transition between OF and IF states. All these functional characteristics are encoded in the free-energy landscape of the protein, or more precisely in the way this landscape is reshaped by one or other ligand, or the lack thereof.

But why does this landscape change? Our simulation data demonstrates that the viability of the alternating-access transition, for a certain binding-site occupancy state, hinges on the energetics of a relatively modest structural change, wherein the network of amino-acids that define the ion-binding sites, as well as the ions themselves, become completely occluded from hydration at either side of the membrane. This open-to-occluded transition, either in the OF or IF states, is only feasible when either 1 Ca^2+^ or 3 Na^+^ ions occupy these sites (alongside several structural water molecules), but not for partial Na^+^ occupancies, or when H^+^ replace Na^+^ or with no ions bound. We expect that other divalent cations that bind to NCX_Mj with significant affinity, such as Cd^2+^ and Mn^2+^ (6), do not permit occlusion either, explaining why they inhibit its activity, rather than being transported (6, 12, 32). It can be concluded, therefore, that the feasibility of this open-to-occluded transition explains the functional specificity of a transporter at the molecular level. This is, in our view, a fundamental insight that is likely to be universally valid. Indeed, distinct conformational changes leading to binding-site occlusion appear to have been captured in experimental structures of other secondary-active transporters, such as Mhp1 (33) vSGLT Faham, 2008 #85; Watanabe, 2010 #86}, LeuT (34–37), LacY (38, 39) and Glt_Ph_ (40). Nonetheless, the actual mechanistic significance of each of these structural observations requires further verification, case by case. To do so, we believe it is imperative to examine these and other transporters through the lens of free-energy landscapes, even though this kind of calculations is technically very challenging and time-consuming. A comprehensive analysis of a symporter would be of particular interest; calculated free-energy landscapes should show occlusion is viable only when the transporter is loaded with all of its substrates, or completely empty. For the latter state, we anticipate the data will show occlusion is feasible because the network of amino-acids involved in substrate recognition can adopt an alternate configuration that maximizes the number of protein-protein interactions, so as to counter the energetic cost of dehydration.

Lastly, it is worth noting that our results confirm that the mechanism of alternating access in the superfamily of cation/Ca^2+^ transporters entails a structural change unlike that observed for other families. This mechanism involves a sliding motion of a two-helix unit across the lipid-bilayer midplane, namely TM1 and TM6, over a distance of about 1 nm. Hinge-like motions within helices TM2 and TM7 facilitate this sliding mechanism, while the remainder of the structure (TM3-TM6, TM8-TM10) is essentially unchanged (**Fig. 6**). Importantly, the configuration of the ion binding sites is also largely unchanged as the transporter switches between outward- and inward-occluded conformations; thus, no channel-like states are ever formed that could allow passive diffusion of the bound ions across the membrane. During the transition between OF and IF conformations, complementary sets of hydrophobic, hydrophilic, and lipid-mediated interactions alternatively form and disrupt at either side of the membrane (**Fig. S6-S8**). Many of these residues have been highlighted by previous mutational analyses for their impact on protein function (T57, F202, G76, D197) or ion recognition (G42, G201, V205, G231, G235) (19, 41). Interestingly, many of these residues are also conserved when the two internal topological repeats are compared (**Fig. S1**), further linking this architectural feature to the alternating-access mechanism. This gradual interconversion between comparable interaction patterns translates into a series of modest free-energy barriers, roughly consistent with measured turnover rate for comparable protein constructs (19). Mammalian orthologs such as cardiac NCX1 are over 1,000 faster than NCX_Mj (29–31), but based on our results, as well other available data (27), it is debatable that this acceleration stems from differences within the ion-binding region; in our view, less conserved regions of the transmembrane domain are the most plausible explanation. Now that the molecular structure of cardiac NCX1 has become available, it will be possible to examine the functional coupling between regulatory and transport domains, through free-energy landscape calculations analogous to those reported here.

## Methods

### Molecular dynamics simulations – general specifications

Conventional and enhanced-sampling MD simulations were carried out using GROMACS2018 or GROMACS 4.5.5 with PLUMED (42–45), at constant temperature (298 K), pressure (1 bar) and periodic-boundary conditions. The simulation systems comprise the protein (with or without ions bound) embedded in a POPC lipid bilayer (of 208 molecules), ∼15,000 water molecules and Cl^-^ counterions, for ∼80,000 atoms in total (**Fig. S3**). The systems were set up using GRIFFIN (46). All ionizable residues in the protein were set to their default state at neutral pH, except D240, which is constitutively protonated (7, 8). To limit the number of possible protein conformations to be sampled and facilitate convergence of the free-energy calculations, we truncated the unstructured intracellular loop between residues A148 and N157, which is not required for substrate recognition or transport function (19). All simulations used the standard CHARMM36m (47) and CHARMM36 (48) force fields for protein and lipids respectively, except for the Lennard-Jones R_min_ parameters for the interactions between carboxylate oxygens and Na^+^ or Ca^2+^, which had been previously optimized against experimental data (7, 8). All the enhanced-sampling simulations were carried out using the bias-exchange Metadynamics method (BE-META) (15).

### Enhanced-sampling simulations of NCX_Mj bound to 3 Na^+^ or 1 Ca^2+^

The enhanced-sampling MD simulations used to calculate the free-energy landscapes shown in **Fig. 2** used as only input the outward-facing, occluded structure of NCX_Mj with 3 Na^+^ ions bound; the protein coordinates are those initially reported (PDB entry 3V5U (6), but the ion-binding site configuration reflects a subsequent correction (7). A total of 18 reaction coordinates were used simultaneously in these calculations; collectively, these coordinates were designed to loosely guide the simulated trajectories to sample the vicinity of an idealized linear path between the OF structure and the repeat-swap (RS) model of the IF state. The functional form of each of these variables is:

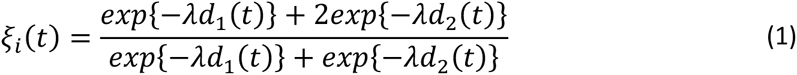

where *d*_1_(*t*), *d*_2_(*t*) quantify the similarity of the configuration at simulation time *t* relative to either of the two reference structures. (Note that neither was assumed to be a free-energy minimum.) The smoothing parameter *λ* is proportional to the inverse of *d*_12_i.e. the similarly between the two end states. The particular metric of similarity used in Eq. 1 is the mean-square difference (MSD) in the configuration of a specific atom selection. Each of the 18 different coordinates *ξ_i_* was defined using a different atom selection. In each case, configurations close to the experimental OF structure correspond to *ξ_i_*∼1, while those close to the RS model correspond to *ξ_i_*∼2. All other configurations result in intermediate values of *ξ_i_*. The specific atom selections that define each of the 18 path-coordinates *ξ_i_* are listed in **Table S1** and graphically depicted in **Fig. S2**. These selections include backbone and sidechains atoms, in both structured and unstructured fragments, both on the intracellular and the extracellular side of the protein; altogether, they describe global and local structural differences between the internal structures of the two topological repeats with the structure of the OF state (**Fig. 1**). Preliminary non-equilibrium simulations of the alternating-access transition using this set of path-coordinates and the minimum-mode Metadynamics method (49) indicated this was a suitable selection.

The BE-META simulations carried out to explore this 18-dimensional space included 32 replicas. In the simplest formulation of the BE-META method, each of the replicas includes a Metadynamics bias that fosters exploration of one reaction coordinate. In previous studies (17, 18) we have observed that the performance of the method is much improved when each replica is designed to explore at least two coordinates, in different combinations (also detailed in **Table S1**). To include a replica for which the sampling is totally unbiased also provides a useful metric of self-consistency. Monte-Carlo exchanges between replicas were attempted every 10-20 ps.

For state with 3 Na^+^ ions bound, we carried out an initial equilibration of 1 μs for each replica; in this time window, all replicas reached uniform sampling in the corresponding reaction-coordinate space, indicating that the bias-potential developed in each replica approximately mirror the corresponding free-energy profile (or potentials-of-mean-force). We also observed that 7 trajectories visited independently an IF state distinct from the RS model. This state was also well populated in the unbiased replica, confirming that it is a low free energy minimum. To initiate the calculations with 1 Ca^2+^ ion bound, we extracted the final configurations from each of the replicas used for the Na^+^-bound state, and in each case, we reconstructed the binding site geometry of Ca^2+^-bound state (**Fig. S10**) through a series of restrained equilibrations totaling 50 ns, while sustaining the biasing potential previously developed in each replica. To fully converge the Metadynamics biasing potentials, we then duplicated the number of replicas to 64 and extended the sampling by 200 ns per replica for the Na^+^ state and by 250 ns per replica for the Ca^2+^ state. The free-energy landscapes shown in **Fig. 2** derive from this 64-replica calculation, for the final 200-ns sampling window. See Supplementary Information for further details.

### Enhanced-sampling simulations of NCX_Mj bound to 2 H^+^, 2 Na^+^ or with no ions bound

Our previous simulation studies of OF NCX_Mj bound to 2 H^+^, 2 Na^+^ or with no ions bound indicated that the alternating-access transition would be limited by the energetics of occlusion of the binding-site region (8), rather than a conformational intermediate between the OF and IF state. Thus, the enhanced-sampling simulations used to derive the free-energy landscapes in **Fig. 5** focused instead on fostering the occlusion or opening of the binding site, by using an alternative set of reaction coordinates. The first type of coordinate quantifies the degree of hydration of each of the ion binding sites. For example, for the central site, known as S_Ca_, this reaction coordinate is:

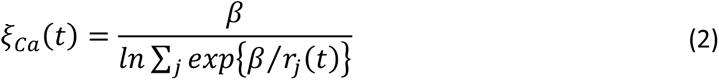

where *r_j_*(*t*) denotes the time-dependent distance between the center of the site and each of the oxygen atoms in any of the water molecules in the system, denoted by the index *j*; β is 10 nm. When the binding site is occupied by an ion, the ion defines its center; when the site is empty, its center is defined by the center of mass of the protein oxygen atoms coordinating the ion when bound. Analogous expressions apply to the other binding sites. The second type of reaction coordinate describes instead changes in the protein structure. For example, in the examination of the IF states, *ξ_TM_*_1,2-7_describes the movement of intracellular portions of TM1 (residues 18-26) and TM2 (residues 45-50) relative to TM7 (residues 211-217), using the same function as that in Eq. 1:

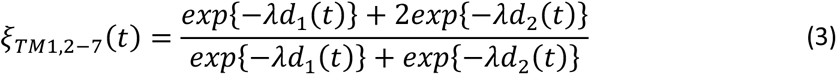

In this case, *d*_1_(*t*) and *d*_2_(*t*) denote the MSD between the configuration at simulation time *t* (defined by Cα atoms of TM1, TM2 and TM7) and either the intracellularly closed or intracellularly open conformation, respectively, and *λ* = 20 nm^-2^. (Reference open and closed conformations were extracted from MD simulations in different ion-occupancy states.) An additional variable of this type describes the relative opening and closing movements of intracellular portions of TM2 (residues 45-50) and TM8 (residues 232-236), and is defined with three reference states:

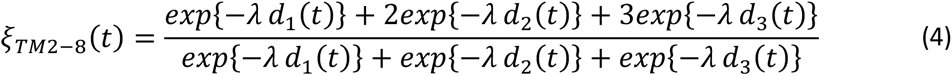

Here, *d*_1_(*t*), *d*_2_(*t*) and *d*_3_(*t*) are the MSD between the current configuration at time *t* and three progressively more open reference structures, and *λ* = 200 nm^-2^.

The BE-META simulations carried out to explore this alternative reaction-coordinate space included 16 replicas; 15 replicas included Metadynamics potentials on different combinations of reaction coordinates, one replica was unbiased. Additional details are provided in **Table S2**. Monte-Carlo exchanges were attempted every 5 ps. The total simulation time was 300 ns for each replica; equilibration of the biasing potentials was achieved after approximately 200 ns, and so only the last 100 ns were used for analysis. See Supplementary Information for further details.

### Derivation of free-energy landscapes from enhanced-sampling trajectories

To translate the BE-META simulation data into 2D free-energy landscapes, we identified two intuitive structural descriptors of intra- and extracellular opening-closing motions. These descriptors are defined by the number of protein-protein contacts between 1) intracellular portions of TM1-TM2 (residues 17, 20, 21, 24, 25, 42, 43, 46, 47, 48, 50) and TM7-TM8 (residues 211, 212, 213, 215, 216, 219, 220, 231, 232, 235, 236); and 2) extracellular portions of TM6-TM7 (residues 176, 179, 180, 183, 184, 201, 202, 205, 206, 207, 209) and TM2-TM3 (residues 52, 53, 54, 56, 57, 60, 61, 72, 73, 76, 77). Note these sets of residues are in pseudo-symmetrical fragments of the two topological repeats. The number of protein-protein contacts for a given residue selection was defined by:

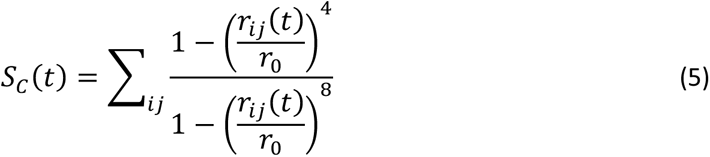

where *r_ij_* denotes the Cα-Cα distance between residue *i* and *j* and *r*_0_, the contact distance, is 0.7 nm.

The free-energy maps shown in Fig. 2A and Fig. 5A reflect a re-weighted histogram as a function of the two descriptors mentioned above. To ensure unbiased sampling, the weight of each simulation frame *i* was set as:

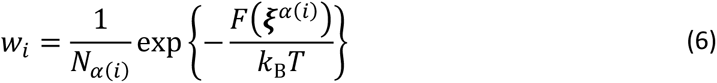

where *F*(***ξ***^*a*(*i*)^) is the free energy as a function of the reaction coordinates used in the BE-META simulations, *⍺*(*i*) is the bin assigned to frame *i* and *N*_1(*i*)_ is the total number of frames in that bin (16). The free energy, *F*(***ξ***^*a*(*i*)^), was derived using the FCAM method (16). Specifically, with either 3 Na^+^ or 1 Ca^2+^ bound, we defined *F*(***ξ***^*a*(*i*)^) as a function of 4 variables, namely *ξ*_5_, *ξ*_1._, *ξ*_15_ and *ξ*_1+_ (**Fig. S2**), and thus this quantity was deduced from the sampling collected in the BE-META replicas in which those variables were biased (**Table S1**). The same replicas were used to derive the free energy maps in **Fig. 2** based on Eq. 6. The minimum free-energy paths in **Fig. 4** were also derived using the FCAM software suite (https://github.com/FCAM-NIH/FCAM), again in the space the 4 reaction coordinates mentioned above.

### Evaluation of simulation results by comparison with experimental HDX data

A methodology was developed to contrast the conformational ensembles obtained with our MD simulations with experimental measurements of hydrogen-deuterium exchange carried out for WT NCX_Mj as well as an elongation mutant 5L6-8. This methodology builds upon a theoretical framework reported in a previous study (22) and is generally applicable to cases where multiple constructs and conformational states are evaluated; it explicitly accounts for the unknown degree of deuterium loss after quenching of the HDX reaction. Further details are provided in Supporting Information.

## Supporting information

Movie of the outward-to-inward conformational change of NCX_Mj bound to one Ca2+ ion. Protein (cartoon), waters (red), and Ca2+ (cyan) are shown.

Movie of the outward-to-inward conformational change of NCX_Mj bound to three Na+ ions. Protein (cartoon), waters (red), and Na+ (green) are shown.

## Acknowledgements

We are thankful to Dr. Emel Ficici for constructing a repeat-swap model of NCX_Mj; to Prof. Daniel Khananshvili and co-workers for providing their published HDX-MS data; and to Dr. Vanessa Leone and Dr. Lucy Forrest for her feedback throughout this research study and their comments on this manuscript. This study was funded by the Division of Intramural Research of the National Heart, Lung and Blood Institute (NHLBI), National Institutes of Health (USA). Most of the computational resources were provided by the NIH high-performance computing facility Biowulf.

## Author contributions

F.M. and J.F.G. conceptualized research. F.M. designed and conducted research. F.M. analyzed and interpreted data. F.M. and J.F.G. wrote the manuscript. J.F.G. supervised and acquired funding and resources for the project.

