## Supporting Figures (Fig. S1-S10), Additional Methods and Supporting Tables (S1,S2) for "Conformational free-energy landscapes of a Na^+^/Ca^2+^ exchanger explain its alternating-access mechanism and functional specificity"

October 19th, 2023

### Supporting Figures

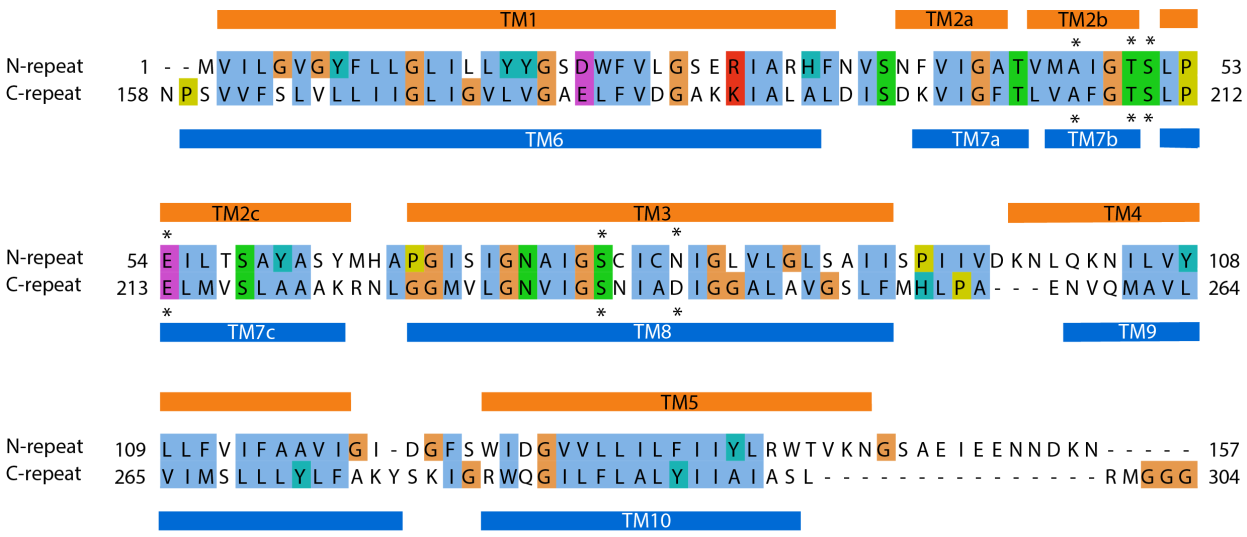

**Figure S1.** Alignment of the amino-acid sequences of the two inverted topological repeats in NCX_Mj. The sequence identity is 32%. The transmembrane regions in each repeat are indicated and color-coded as in Fig. 1. Residues directly involved in ion-binding are marked with asterisks. The repeat-swap model shown in Fig. 1 was constructed using this alignment.

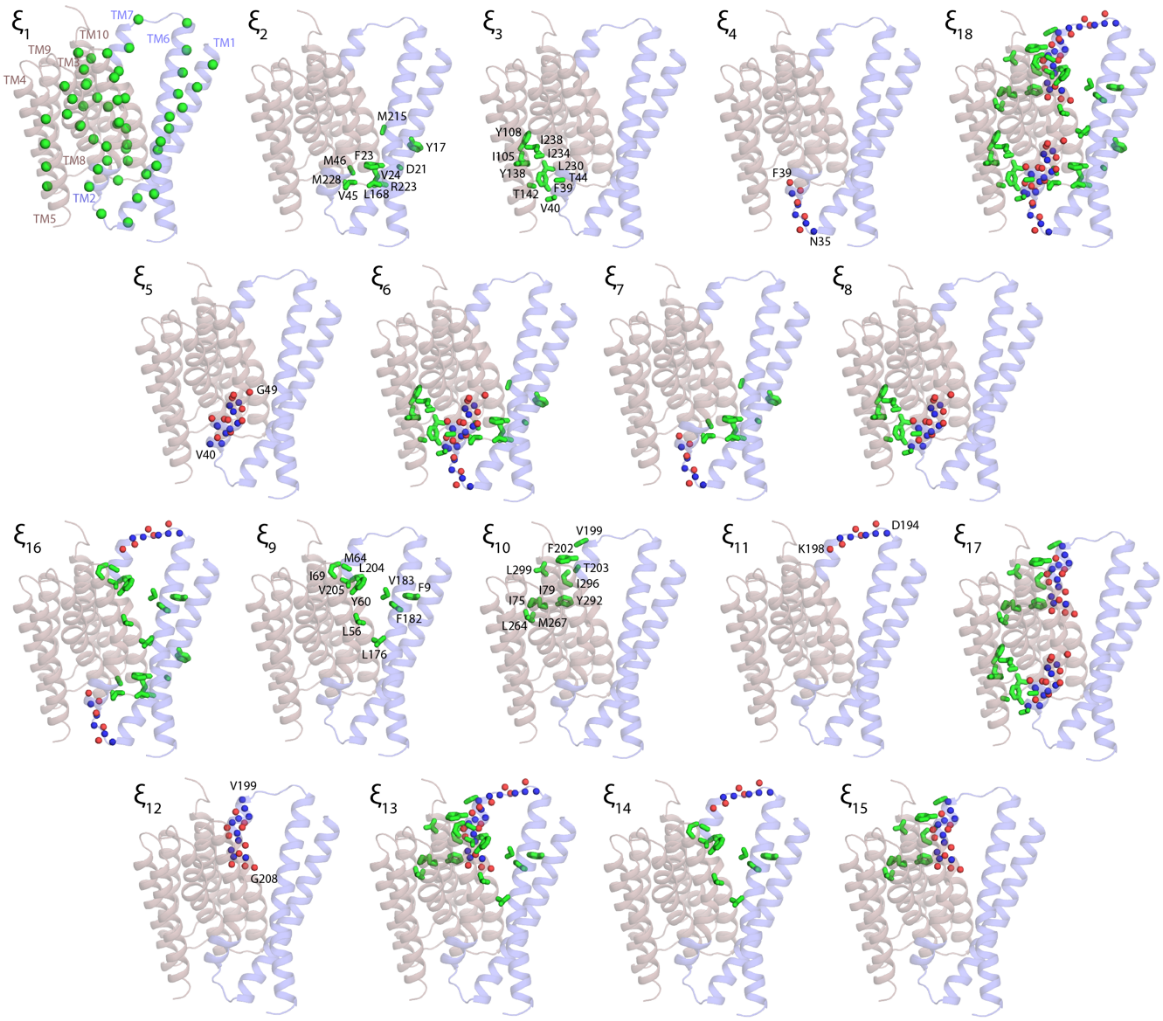


**Figure S2**. Atom selections used to define the path-collective variables employed in the bias-exchange Metadynamics simulations of the alternating-access transition (see Methods). The specific reaction coordinate associated with each selection is indicated (see also Table S1). The selections include Cα atoms (green spheres), backbone N and O atoms (blue and red spheres), and sidechain carbon atoms (green sticks). Note these selections are approximately symmetric, i.e. they include equivalent residues in both topological repeats.

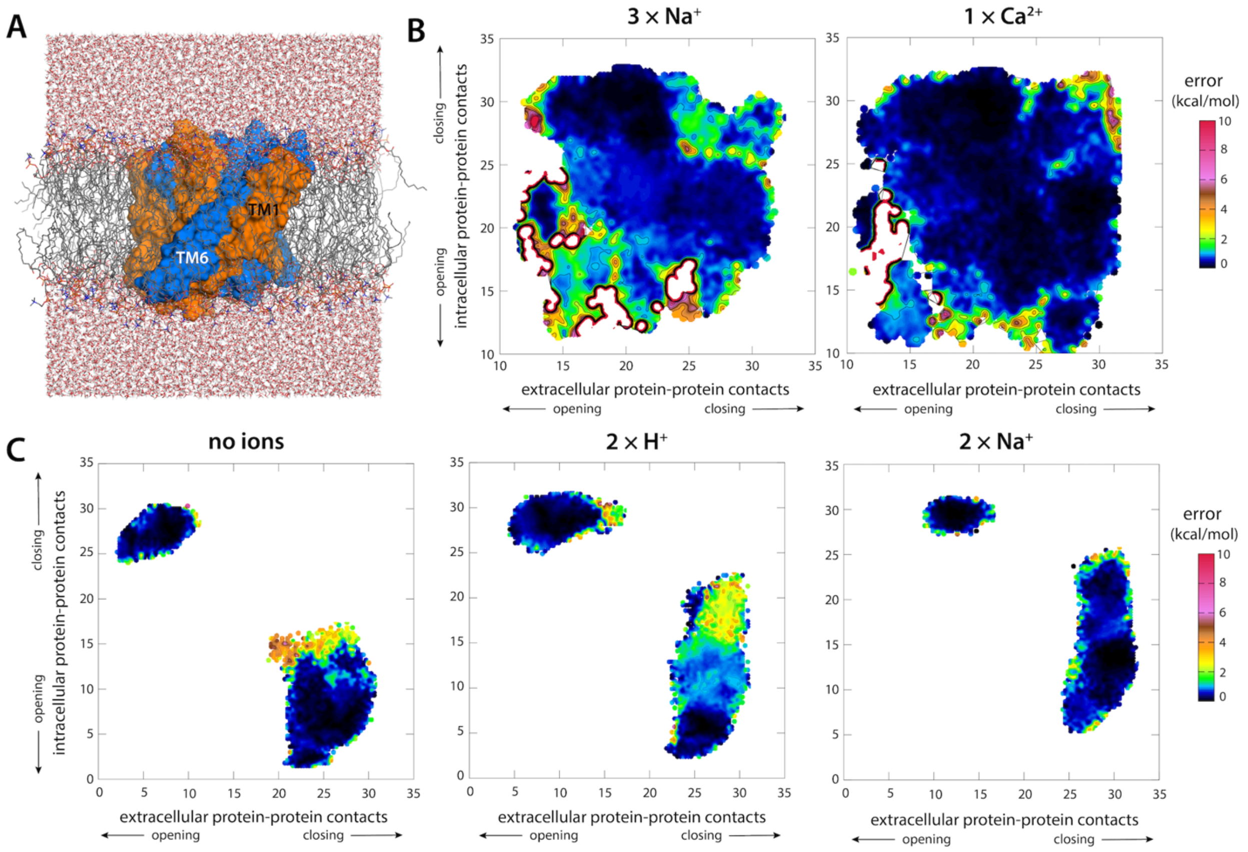


**Figure S3**. Molecular simulation system and error estimates. (**A**) All-atom simulation system used to calculate the free-energy landscapes shown in Fig. 2 and Fig. 5. The transporter (orange/blue) is embedded in a hydrated POPC lipid bilayer; electrolyte ions are omitted for clarity. (**B**) Estimated error of the free-energy maps shown in Fig. 2, based on block analysis. (**C**) Estimated error of the free-energy maps shown in Fig. 5, also from block analysis.

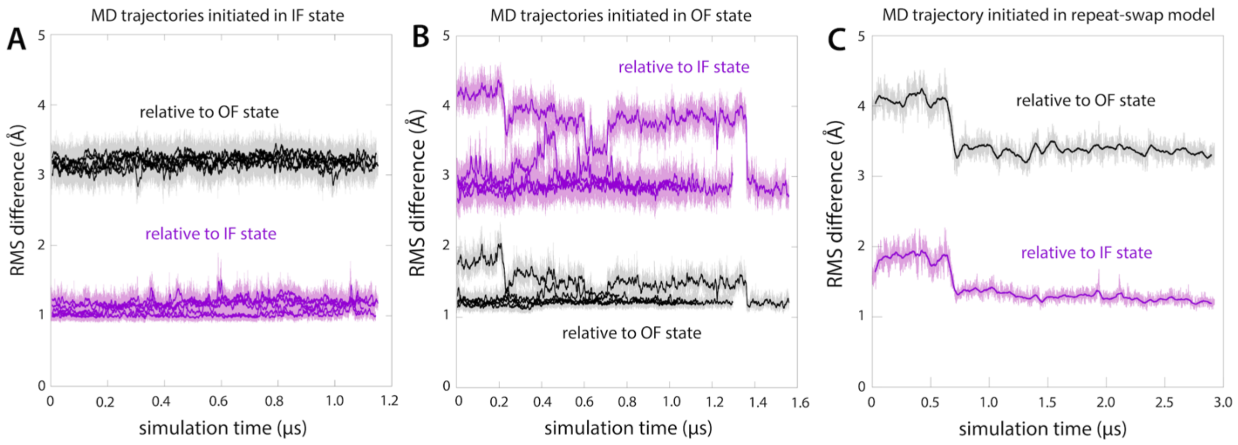


**Figure S4**. Conventional MD simulations of NCX_Mj bound to 3 $Na^+$ ions. (**A**) Trajectories initiated in the inward-facing (IF) occluded state identified in the free-energy landscape shown in Fig. 2. The plot quantifies the average root-mean-square (RMS) difference between the $C_\alpha$ trace at a given simulation time and the $C_\alpha$ traces in an ensemble of structures extracted from each of the two main free-energy basins in that landscape (within 2 kcal/mol of minimum), i.e. the original IF state, and the outward-facing (OF) occluded state. Unstructured loop regions were omitted in this analysis. (**B**) Same as (A), for trajectories initiated instead in the OF occluded state. (**C**) Same as (A), for a trajectory initiated in the repeat-swap model shown in Fig. 1.

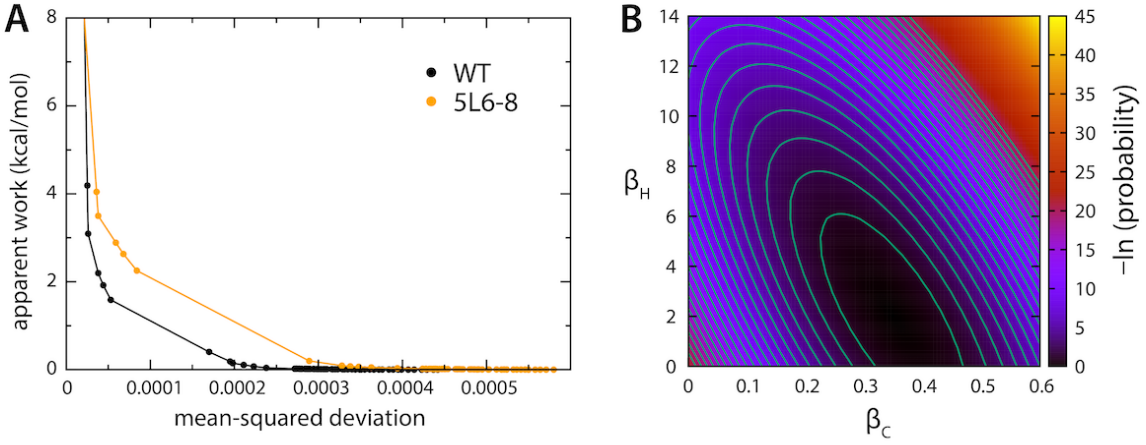


**Figure S5**. Evaluation of the predicted conformational ensembles for OF and IF states, in reference to HDX-MS data for two forms of NCX_Mj with different propensities for the OF and IF states. See also Fig. 3C. (**A**) The plot quantifies the degree to which a conformational ensemble that initially consists of equally weighted populations OF and IF structures (extracted from simulation) must be re-reweighted to produce optimal agreement with the experimental data obtained for each protein construct (see Additional Methods). The degree of reweighting is quantified by an 'apparent work', and the agreement with experiment by the mean-squared deviation between calculated and measured deuteration fractions for a collection of protein fragments. (**B**) Prior probability distribution of the model parameters used to predict hydrogen-deuterium exchange protection factors based on the forward model of Best and Vendruscolo (1).

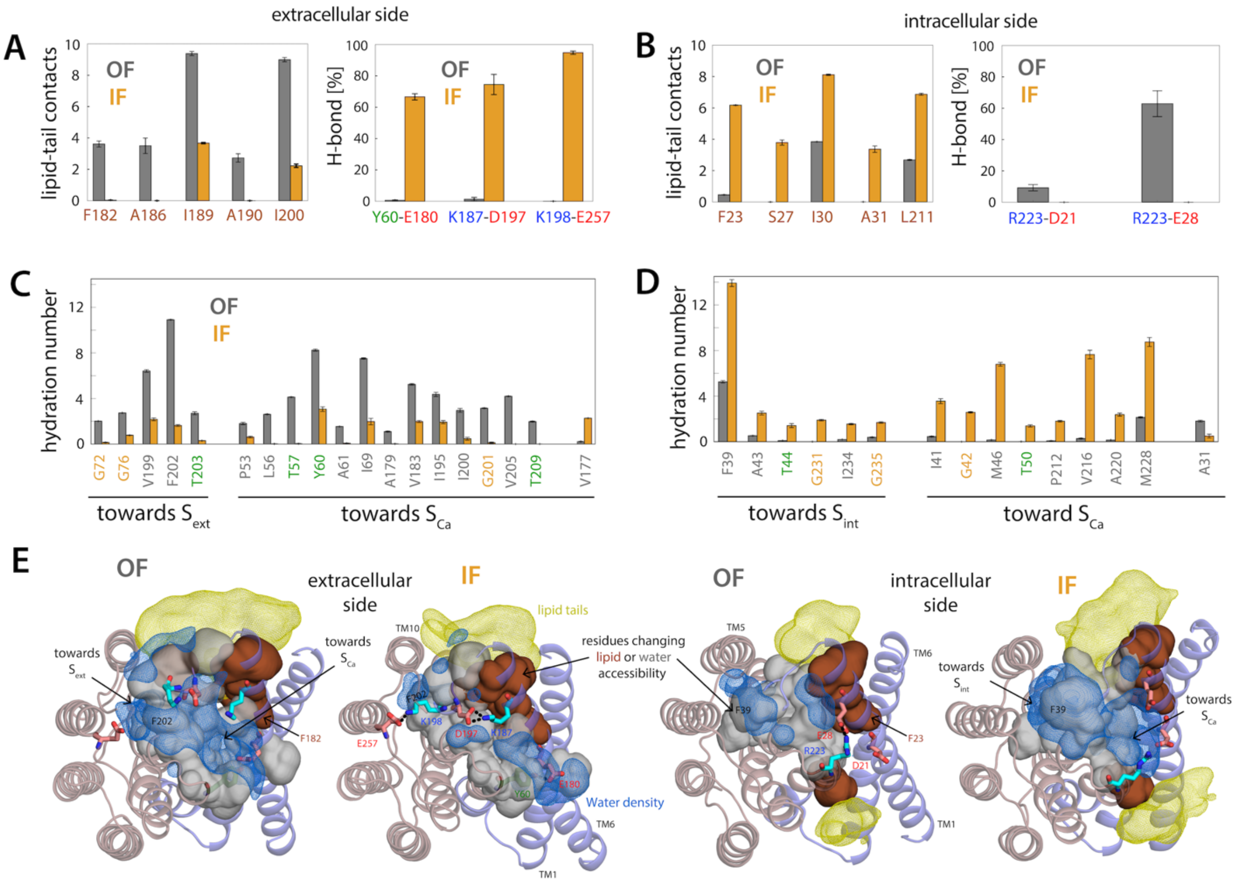


**Figure S6**. Changing interaction patterns in the alternating-access mechanism of NCX_Mj. Reported quantities are calculated for ensembles of either OF or IF structures, with 3 $Na^+$ ions bound, extracted from the free-energy basins revealed in the landscapes shown in Fig. 2. (**A, B**) The charts enumerate the set of residues in the extracellular or intracellular side of the protein, respectively, that show a significant change in hydrophobic lipid-tail contacts or protein-protein H-bonding and salt-bridges when OF and IF stats are compared. (**C, D**) The charts enumerate the set of residues in the extracellular or intracellular side of the protein, respectively, that show a change in hydration number of 2-fold or more; the hydration number was defined using a cutoff distance of 4.2 Å. (**C**) Representative structures of the OF and IF ensembles illustrating the changes in interaction patterns quantified in panels (A,B) and (C,D). The residues that exhibit changes in water accessibility and lipid contacts are shown as grey and brown volumes, respectively, with water and lipid molecules in their vicinity shown with density isosurfaces (blue and yellow mesh, respectively). Residues that exhibit changes in protein-protein H-bonding and salt bridges are shown as sticks. The number of lipid-tail and H-bond contacts between two groups of atoms in a given simulation snapshot was calculated using distance cut-off values of 5.2 and 3.2 Å, respectively. Hydrogen atoms were excluded from this analysis.

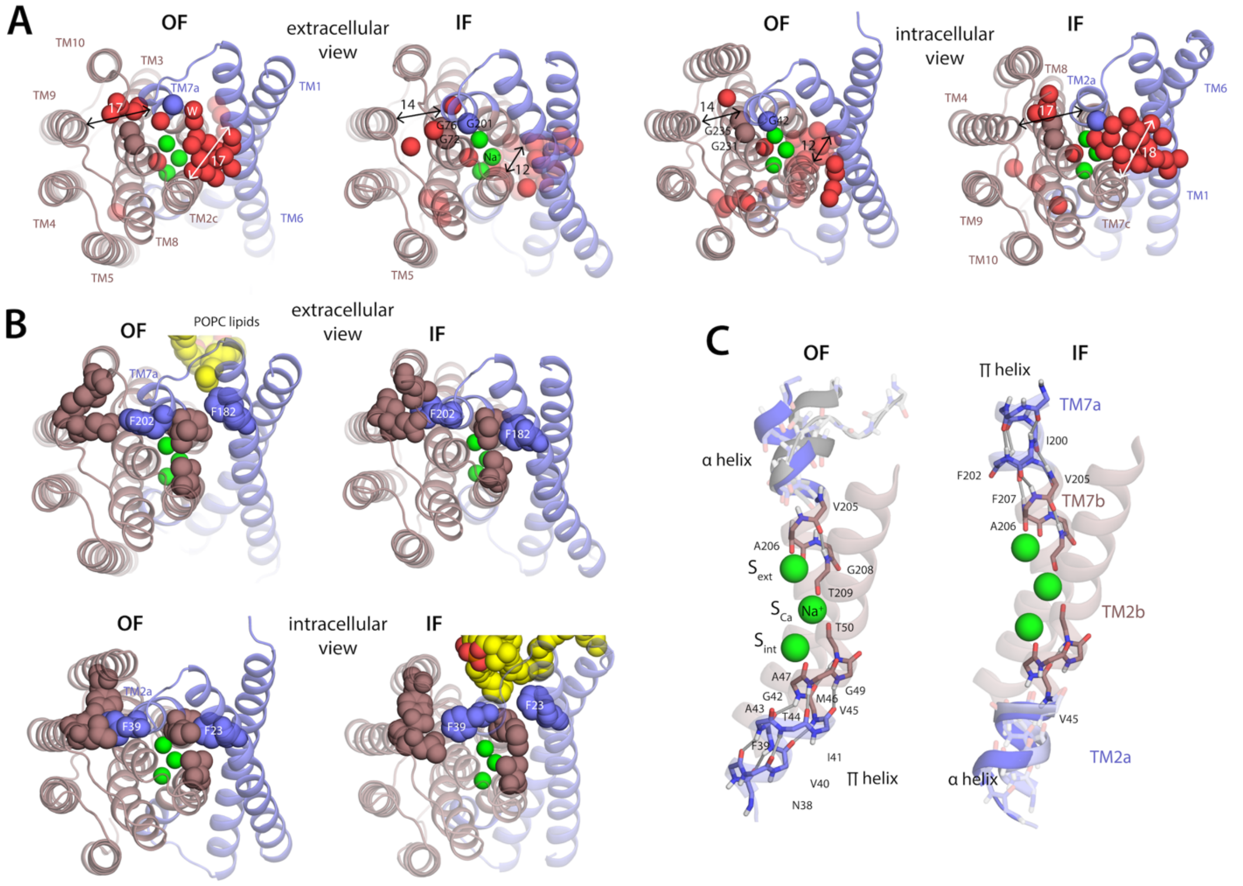


**Figure S7**. Changing interaction patterns in the alternating-access mechanism of NCX_Mj. (**A**) Extracellular and intracellular views of representative OF and IF conformations, with 3 $Na^+$ ions bound (green spheres), highlighting water molecules within 10 Å of the ion-binding sites (red spheres). The approximate distances between TM helices involved in the opening and closing of intra- and extracellular pathways are indicated in Å. (**B**) Same views as panel (A), showing phenylalanine sidechains (spheres) that occupy or become released from hydrophobic pockets during the alternating access transition (F182 and F202 on the extracellular side, and F23 and F39 on the intracellular side). Protein residues that are in contact with those phenylalanine sidechains when buried in a hydrophobic pocket are also highlighted, as are lipid molecules in contact with either F23 or F182 when disengaged (yellow). (**C**) Close-up of the backbone of N-terminal portions of TM2 (TM2a-TM2b) and TM7 (TM7a-TM7b) in the OF and IF state, highlighting selected intramolecular H-bonds (gray lines).

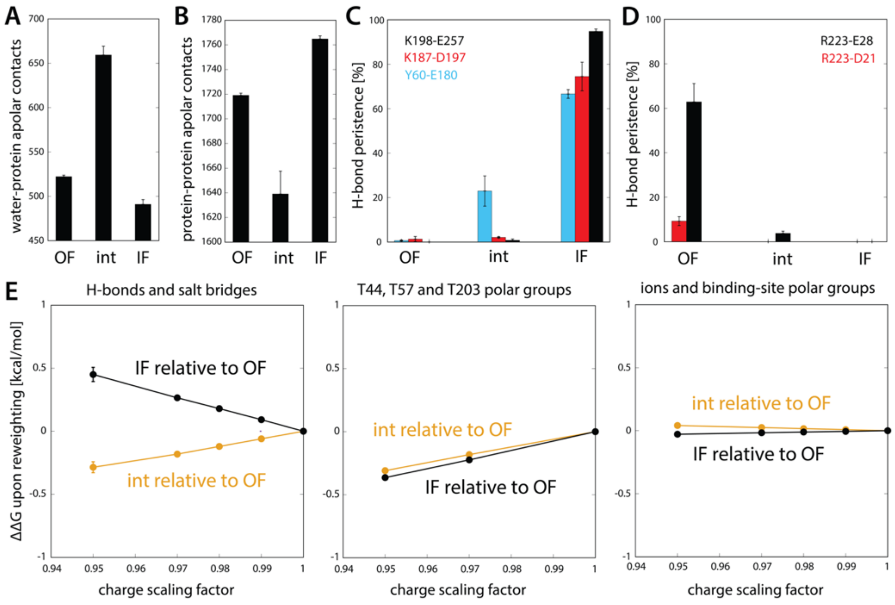


**Figure S8**. Global changes in hydrophobic, H-bond and electrostatic interactions alternating-access mechanism of NCX_Mj. Reported quantities are calculated for ensembles of either OF or IF structures, with 3 $Na^+$ ions bound, extracted from the free-energy basins revealed in the landscapes shown in Fig. 2. Data is also shown for an ensemble of intermediate configurations. (**A**) Number of contacts between water and protein apolar groups in each state, calculated as in Fig. S6, for $r_0$ = 4.2 Å .The residues selected are those that change significantly their accessibility to water during the alternating-access transition, namely: 24, 31, 38, 39, 41, 42, 43, 44, 46, 50, 53, 56, 57, 60, 61, 66, 68, 69, 72, 76, 138, 177, 179, 183, 195, 198, 199, 200, 201, 202, 203, 205, 209, 212, 216, 220, 225, 228, 231, 234, 235, and 260. (**B**) Number of contacts between the apolar portion of the previously mentioned protein residues and the apolar portion of all other protein residues, for each state. The number of contacts between two groups of atoms indexed by $i$ and $j$, for a given simulation snapshot at time t, was calculated as $S(t) = \sum_{ij}[1-\left(r_{ij}(t)/r_0\right)]^8/[1-\left(r_{ij}(t)/r_0\right)]^{16}$, where $r_{ij}$ is the distance between any pair of atoms and $r_0$ is a constant that depends on the type of interaction; the values used for water-protein and protein-protein contacts were 4.2 and 5.2 Å, respectively. (**C**) Persistence of H-bonding interactions between residues on the extracellular side of the protein, for each state; only interactions that change significantly during the transition are examined. (**D**) Same as (C), for residues on the intracellular side. (**E**) Analysis of the contribution of electrostatic interactions to the free-energy difference between the OF state and either the IF or the intermediate state, evaluated after scaling down the charges of selected atom groups (via a multiplication factor). To quantify the contribution of H-bonds and salt bridges, we scaled down the atomic charges in the polar groups of residues D21, E28, Y60, E180, K187, D197, K198, R223 and E257 (specifically, to the atoms in each of the 'charge groups' bearing a either a neutral or finite integer charge e.g. the $-CH_2-COO^-$ group of glutamate or aspartate). To assess the contribution of polar residues changing accessibility during the transition, we scaled down the charges of the polar groups of T44, T57 and T203. The contribution of the binding site was evaluated by scaling down the charges of the polar groups of residues coordinating the bound $Na^+$ and water molecules, as well the ions and water molecules. To estimate the change in these free-energy differences, we resampled the BE-META trajectories (with 3 $Na^+$ bound, for the replicas with bias on $\xi_6$, $\xi_{13}$, $\xi_{16}$ or $\xi_{17}$; see Table S1), introducing a reweighing factor for each configuration given by $W_i = exp\ \{-(U_i^C - U_i)/k_BT$, where the term $(U_i^C - U_i)$ is the difference between the potential-energy functions with modified and conventional charges, respectively.

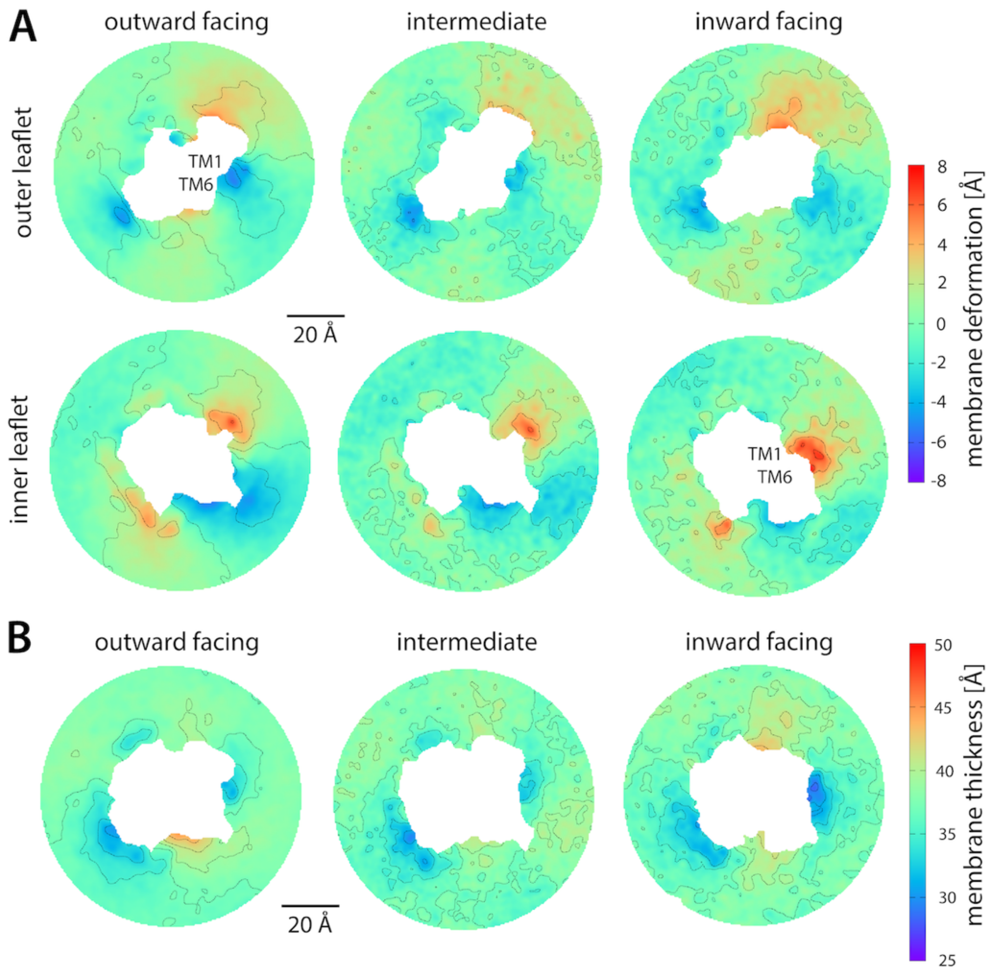


**Figure S9.** Changes in membrane shape and thickness during the alternating access transition. (**A**) Deformation of each of the lipid bilayer leaflets relative to a flat shape, for the OF, intermediate and IF states. To quantify this deformation, we evaluated the mean Z-coordinate of the ester layer across the XY plane relative to the mean Z-coordinate of both layers, in the bulk region (at a distance larger that 45 Å from the ion-binding sites center, where the membrane is approximately flat). (**B**) Hydrophobic thickness of the lipid bilayer in the OF, intermediate and IF states. The thickness was evaluated as the difference between the mean Z-coordinates of the outer and inner ester layers. Contours are shown in intervals of 2 Å. All maps were calculated after first aligning the protein onto a reference frame, using the portion of the structure that is invariant when OF and IF states are compared (Fig. 2). To prevent tilting of the membrane during this process, this alignment was two-dimensional (i.e. only X and Y components of the atomic coordinates are considered). After structural alignment we calculated for each state a three-dimensional occupancy map for the ester oxygen atoms of all POPC lipids in either the outer or the inner leaflet, using the Volmap tool of VMD {Humphrey, 1996 #124}. The mean Z-coordinate was calculated as the weighted average of the Z-coordinate for each value of X and Y, with a weight provided by the occupancy at that position. Values of X and Y featuring less than 1% of the overall occupancy are excluded (and correspond to the region occupied by the protein in the center of each map).

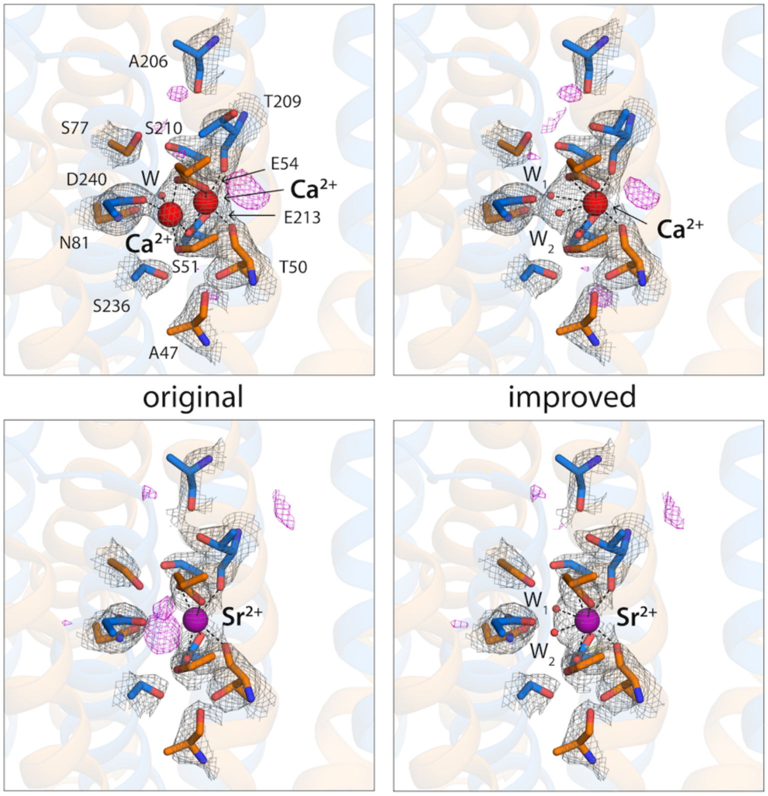


**Figure S10**. Improved molecular refinement of the $Ca^{2+}$ and $Sr^{2+}$ coordination geometries in outward-facing NCX_Mj based on previously published crystallographic data (2). The original models (PDB entries 5HXR and 5HXS), shown in the left, are contrasted with our proposed interpretation, shown on the right. 2Fo−Fc maps contoured at 1σ is shown as a grey mesh and Fo−Fc omit maps at 3σ is shown as in magenta. See Supplementary Methods for further details.

### Additional Methods

*Construction and simulation of repeat-swap model of NCX_Mj*

To construct the repeat-swap model of NCX_Mj shown in Fig. 1, we first generated a reference sequence by swapping the residues of the two repeats, with the specific sequence alignment provided in Fig. S1, as described elsewhere (3). We then generated 250 models with MODELLER v9.15 (4) using as template the OF X-ray structure of NCX_Mj bound to 3 $Na^+$ ions (5). To preserve the $C_2$ pseudo-symmetry of the binding site region with respect to an axis parallel to the membrane plane, we restrained the distances between the 3 $Na^+$ ions and their coordinating oxygens as well as the oxygen-oxygen distances. The lowest-energy model was selected for further refinement. First, sidechain rotamers were optimized using SCWRL4 (6). Then, the model was energy-minimized according to the CHARMM27/CMAP forcefield (7, 8) and embedded in a solvated POPC bilayer using GRIFFIN (9). An MD simulation of 800 ns was then carried out to equilibrate the model at constant temperature (298 K), pressure (1 atm) and membrane surface area (~69 Å$^2$ per lipid). This simulation was carried out with NAMD 2.12(10) and the same force field. This initial equilibration involved a series of stages in which first position restraints and then RMSD restraints are gradually removed on protein sidechain and backbone atoms. Thereafter we performed additional 3 μs MD simulation without any restraint (see Fig. S4 for results).

*Improved refinement of crystallographic data for NCX_Mj bound to $Ca^{2+}$ and $Sr^{2+}$*

A statistical survey of the $Ca^{2+}$-binding sites in the PDB (2) suggests that when $Ca^{2+}$ is bound to two carboxylate groups in a bidentate mode, the latter ion has a total coordination of eight. To fulfill this coordination geometry, in our previous simulation study of NCX_Mj, based on the OF X-ray structure (2), we placed $Ca^{2+}$ in the $S_{Ca}$ site and introduced two water molecules coordinating $Ca^{2+}$ at near the $S_{mid}$ site. This ion coordination mode has also been used in this study. By contrast, in the original interpretation of the crystallographic data, $Ca^{2+}$ was assumed to partially occupy $S_{Ca}$ and $S_{mid}$ sites, assisted by one water molecule (2) (Fig. S10). To validate or refute our previous interpretation, we carried out a new refinement of NCX_Mj bound to $Ca^{2+}$ based on the electron-density map available in the Protein Data Bank (entry 5HXR). Model building was completed with COOT (11), and structure refinement was carried out with PHENIX (12). The results are shown in Fig. S10. It is clear that the geometry we propose for the $Ca^{2+}$ state is just as compatible with the electron-density data as the original model; it is however more plausible, statistically speaking, and is also supported by recent ATR-FTIR experiments (13), which show that $Ca^{2+}$ does not reside near residue D240 at the $S_{mid}$ site. To further support the proposed coordination geometry, we also carried out a new refinement of crystallographic data obtained for $Sr^{2+}$(2), which is also transported by NCX_Mj (14). This analysis clearly reveals that in the published structural model (entry 5HXS) the coordination of the $Sr^{2+}$ ion is incomplete since additional coordinating atoms ought to be present near the $S_{mid}$ site (Fig. S10). Indeed, a new refinement adding two coordinating water molecules in the defective region significantly improves the agreement with the crystallographic data (Fig. S10). The conclusion from this analysis is that both $Ca^{2+}$ and $Sr^{2+}$ bind only at the $S_{Ca}$ site and in a nearly identical mode, which includes two coordinating water molecules at the $S_{mid}$ site.

*Restraining potentials used in the bias-exchange Metadynamics calculations*

In the bias-exchange Metadynamics (BE-META) simulations of NCX_Mj with 2 or 3 $Na^{+}$ ions bound, we applied restraining potentials on a coordination variable to preclude $Na^{+}$ dissociation (even though spontaneous dissociation was not observed in unbiased simulations). The coordination variable was defined as:

$$\xi_C(t) = \sum_k \frac{1-\left(\frac{r_k}{r_0}\right)^8}{1-\left(\frac{r_k}{r_0}\right)^{10}} \qquad \text{(S1)},$$

where $r_k$ is the distance between ions and coordinating oxygen atoms and $r_0 = 0.24$ nm. For this variable we set lower values of $\xi_0 = 4.3$ for $Na^{+}$ ions bound to $S_{int}$ and $S_{ext}$ sites (coordinated to 5 oxygens) and $\xi_0 = 4.75$ for the ion bound to site $S_{Ca}$ (coordinated to 6 oxygens; Fig. 1C), using a potential of the form $k_C(\xi_C(t) - \xi_0)^4$, applied when $\xi_C(t) < \xi_0$ and for which $k_C = 1000$ kJ/mol. The water molecule in the binding site (Fig. 2C) was confined with two half harmonic potentials, $k_d(r(t) - r_0)^2$, on distances, $r(t)$, between the water oxygen and the $C_\gamma$ atoms of N81 and D240 respectively. This potential was set with $k_d = 10000$ kJ/(mol nm$^2$) and applied when the above mentioned distances are larger than $r_0 = 0.46$ nm.

A similar scheme was used in the BE-METAD simulations with $Ca^{2+}$ bound, but only for the $S_{Ca}$ site. This potential was applied on the coordination variable defined by Eq. S1 in which $r_k$ is the distance between $Ca^{2+}$ and a coordinating oxygen atom, and $r_0 = 0.3$ nm. For this variable we set a lower value of $\xi_0 = 7.4$ using a potential of the form $k_C(\xi_C(t) - \xi_0)^2$, applied when $\xi_C(t) < \xi_0$ and for which $k_C = 400$ kJ/mol. The two water molecules at the $S_{mid}$ site (Fig. 2C) were also confined with a half harmonic potential, $k_d(r(t) - r_0)^4$, on the distances ($r$) between the $C_\gamma$ atoms of either N81 or D240 residues with the oxygen atom of the adjacent water molecule. For this potential $r_0 = 0.41$ nm and $k_d = 10{,}000{,}000$ kJ/(mol nm$^4$). To prevent spurious water penetration and facilitate convergence, we also applied a boundary potential on the coordination variable defined by Eq. S1 in which $r_k$ is the distance between $Ca^{2+}$ and all oxygen atoms of water molecules (excluding the those coordinating $Ca^{2+}$), and $r_0 = 0.3$ nm. Distances, $r_k$, beyond 0.5 nm were excluded. For this variable we set an upper value of $\xi_0 = 0.5$ using a potential of the form $k_C(\xi_C(t) - \xi_0)^2$, applied when $\xi_C(t) > \xi_0$ and for which $k_C = 400$ kJ/mol. To prevent unfolding of binding site portions of TM2 and TM7 we also applied harmonic distance restraints ( $k_d(r(t) - r_0)^2$, with $r_0 = 0.41$ nm and $k_d = 10000$ kJ mol$^{-1}$ nm$^{-4}$) between: 1) carbonyl oxygen of A206 and side chain oxygen of S210; 2) carbonyl oxygen of A206 and amide nitrogen of S210; 3) carbonyl oxygen of V205 and amide nitrogen of T209; 4) homologous distance pairs based on inverted symmetry (A47-S51, M46-T50).

*Maximum-entropy reweighting of the simulated ensembles based on HDX-MS data*

In its original formulation, the HDXer (15) methods assumes that the input HDX-MS datasets are corrected for deuterium back-exchange. While this is an ideal situation, not all datasets can be produced with this additional layer of information. In particular the HDX-MS data for NCX_Mj do not include this correction (personal communication from Prof. Khananshvili). Therefore, it was necessary to develop an additional strategy that allows for reweighting ensembles on data that have not been corrected for back-exchange levels.

This new scheme assumes that two HDX-MS datasets are available, for which uncorrected deuterium fractions (with values ranging from 0 to 1) of peptide *j* measured at time *t* are denoted as $D_{j,t}^{exp1}$ and $D_{j,t}^{exp2}$ respectively. These datasets refer to experiments carried out under two different biochemical/physiological conditions (e.g., the presence or absence of ligands) or for two different mutants of the same protein. All HDX experiments are assumed to be performed with the same protocol so that the back-exchange contribution to a given peptide can be reasonably assumed to be the same across the two datasets. In case those datasets correspond to measurements carried out on two protein variants, only identical peptides must be considered for the analysis (i.e., peptides containing sequence variations should be excluded).

A key approximation in this approach is that back-exchange is accounted for by normalization of the deuterated fraction of a given peptide by a factor, $c_j$, which retains approximately the same value for different timepoints (16, 17) and, as previously mentioned, also across the two different HDX datasets. Provided a fully deuterated control were available, the factor $c_j$ could be experimentally determined using the relation: $c_j = D_{\%}\, n_j/(m_{100} - m_0)$, where $m_{100}, m_0$ are the centroid masses of the fully-deuterated and non-deuterated peptide respectively, $D_{\%}$ is the D/H fraction in the buffer and $n_j$ is the associated number of exchangeable amide hydrogens. However, in this case we assume that the experimental data are not corrected for back-exchange. Hence, the term $c_j$ is unknown and therefore is set as a free parameter during the reweighting procedure. The implication is that, rather than the absolute experimental deuteration levels, the target of the ensemble reweighting is primarily determined by the variations between the deuterium fractions of the two datasets. Hence, the sensitivity of the reweighting procedure will depend on the magnitude of these variations (in addition to the number of data points).

The proposed scheme involves two concurrent ensemble reweighting calculations, each targeting one of the HDX-MS datasets, and in which the factors $c_j$ are determined at each reweighting step by best fitting to both datasets. The overall likelihood optimization function is given by the sum of the two individual contributions for each dataset (1 and 2):

$$L = \frac{W_{app}^{1}}{k_B T} + \frac{W_{app}^{2}}{k_B T} - \gamma ln P_{err} \qquad \text{(S2),}$$

where $W_{app}^{k=1,2}$ is the apparent work required to reweight the ensemble and $P_{err}$ is the error distribution of the experimental data, including both HDX datasets.

To compute the work requires a model of the exchange for a given frame in the ensemble. As in our previous work, we use the exchange model of Best and Vendruscolo (1):

$$ln P_i^{simk} = \langle \beta_{\mathrm{H}} N_{\mathrm{H},i} + \beta_{\mathrm{C}} N_{\mathrm{C},i} \rangle_k \qquad \text{(S3),}$$

in which $P_i^{simk}$ is the predicted protection factor of residue *i* from reweighted ensemble *k*, $N_{\mathrm{H},i}$ and $N_{\mathrm{C},i}$ are the number of H-bonds and heavy-atom contacts of the amide group of residue *i,* respectively, and $\langle \dots \rangle_k$ denotes a mean value over the reweighted ensemble *k*. Then, based on the maximum entropy principle, the apparent work can be expressed as (15):

$$\begin{aligned} W_{app}^{k=1,2}/k_B T = &\sum\nolimits_i \left( {}^k\lambda_i^C \langle N_i^C \rangle_k + {}^k\lambda_i^H \langle N_i^H \rangle_k \right) \\ &- ln \int exp\left\{ -\beta U_{MD}(\mathbf{X}_{\boldsymbol{k}}) + \sum\nolimits_i \left( {}^k\lambda_i^C N_i^C(\mathrm{X}_k) + {}^k\lambda_i^H N_i^H(\mathrm{X}_k) \right) \right\} d\mathbf{X}_{\boldsymbol{k}} \end{aligned} \qquad \text{(S4),}$$

where $N_i^H(\mathrm{X}_k)$ and $N_i^C(\mathrm{X}_k)$ are the number of H-bonds and heavy atom contacts, respectively, for the amide of residue *i*, at a molecular configuration, $\mathrm{X}_k$. We assume that the error distribution, $P_{err}$, is Gaussian:

$$P_{err} \propto exp\left\{ -\sum\nolimits_k \sum\nolimits_t \sum\nolimits_j \left( D_{j,t}^{simk} - c_j D_{j,t}^{expk} \right)^2 / 2\eta_D^2 \right\} \qquad \text{(S5),}$$

The normalization parameters, $c_j$, are determined by minimizing at each step the mean square deviation between simulated and experimental deuteration fractions (i.e. the exponent in Eq. S5):

$$\begin{gathered} \frac{\partial \left( \sum_t \sum_{k=1,2} \left( D_{j,t}^{simk} - c_j D_{j,t}^{expk} \right)^2 + A_j(c_j - 1) + B_j \left( c_j - \frac{1}{D_{j,t}^{expk}} \right) \right)}{\partial c_j} = 0 \\ c_j \geq 1 \\ c_j \leq \frac{1}{D_{j,t}^{expk}} \end{gathered} \qquad \text{(S6),}$$

where the constants $A_j$ and $B_j$ are (Lagrange multipliers) set to satisfy the boundary conditions defined by the two inequalities in Eq. S6. The condition $c_j \geq 1$ stems from the consideration that, owing to the back-exchange, $D_{j,t}^{expk}$ is a lower bound of the correct deuteration level. The other boundary constraint ($c_j D_{j,t}^{expk} \leq 1$) instead ensures that the corrected deuteration fraction has an upper boundary of 1. Solving Eq. S6 leads to the following expression for the normalization terms:

$$c_j = \frac{A_j + B_j + \sum_{k=1,2}\sum_t D_{j,t}^{simk} D_{j,t}^{expk}}{\sum_{k=1,2}\sum_t \left(D_{j,t}^{expk}\right)^2} \quad \text{(S7),}$$

where $A_j = B_j = 0$ in the case that all boundary constraints are satisfied.

As in our previous work (15), the terms ${}^{k}\lambda_i^H$ and ${}^{k}\lambda_i^C$ are iteratively determined in order to minimize the likelihood function of Eq. S2:

$${}^{k}\lambda_i^{C/H}[n+1] = {}^{k}\lambda_i^{H/C}[n](1-\epsilon) + \epsilon\gamma\beta_{C/H}\frac{\partial lnP_{err}}{\partial lnP_i^{simk}} \quad \text{(S8),}$$

where $n$ denotes the iteration step, $\gamma$ is a parameter that sets the experimental uncertainty and $\epsilon$ is a suitably chosen update rate.

*Optional optimization of the model parameters*

In the previous section we assumed that the model parameters, $\beta_H$ and $\beta_C$, have a fixed value, usually set to that derived by Best and Vendruscolo, which best fit a set of HDX measurements ($\beta_H = 2.0$, $\beta_C = 0.35$) (1). To reduce model bias, we set $\beta_H$ and $\beta_C$ as optimizable parameters during reweighting, which are sampled using a Monte Carlo algorithm and averaged out during reweighting. The parameter sampling stems from choosing an error probability distribution in which $\beta_H$ and $\beta_C$ are integrated out:

$$P_{err} = \int exp\left\{-\sum_k\sum_t\sum_j \left(D_{j,t}^{simk} - c_j D_{j,t}^{expk}\right)^2 / 2\eta_D^2\right\} P(\beta_H\beta_C)d\beta_H d\beta_C \quad \text{(S9),}$$

where $P(\beta_H\beta_C)$ is the prior error distribution of the model parameters. Applying a prior distribution can be useful for sparse and noisy experimental data, as it restricts the sampling to a reasonable range of values. Here, $P(\beta_H\beta_C)$ was set to mimic the trend of the square residuals as a function of $\beta_H$ and $\beta_C$ (Fig. S5B) that was reported by Best and Vendruscolo (1). Specifically, those square residuals describe the discrepancy between predicted and experimental protection factors for a set of proteins. Based on Eq. S9, the update rule for ${}^{k}\lambda_i^{C/H}$ becomes:

$${}^{k}\lambda_i^{C/H}[n+1] = {}^{k}\lambda_i^{H/C}[n](1-\epsilon) + \epsilon\gamma_D \langle\beta_{C/H}\frac{\partial ln\acute{P}_{err}(\beta_H,\beta_C)}{\partial lnP_i^{simk}}\rangle_{\beta_C,\beta_H} \quad \text{(S10),}$$

where $\langle\ldots\rangle_{\beta_C,\beta_H}$ denotes an average performed over $\beta_C,\beta_H$ parameter space, with probability $\acute{P}_{err}(\beta_H,\beta_C) = exp\left\{-\sum_k\sum_t\sum_j\left(D_{j,t}^{simk} - c_j D_{j,t}^{expk}\right)^2/2\eta_D^2\right\}P(\beta_H\beta_C)$. The term $\eta_D^2$ was set as the minimum value (as a function of $\beta_C,\beta_H$) of the mean square deviation, $MSD = \sum_k\sum_t\sum_j\left(D_{j,t}^{simk} - c_j D_{j,t}^{expk}\right)^2/N$, before reweighting, in which $N$ is the number of data points.

The average in Eq. S10 is performed using Monte Carlo with acceptance probability provided by $\acute{P}_{err}(\beta_H, \beta_C)$.

*Additional details on the application of the reweighting methodology to NCX_Mj*

The original HDXer formulation (15) is based on reweighting single protein configurations, which could lead to different weights for very similar structures, for example due to a small structural variation that is sufficient to break a few amide H-bonds. To prevent this, instead of single configurations we reweight free-energy bins. The free-energy of bin $\alpha$ corrected according to the maximum entropy principle is defined as:

$$F_{corr}(\boldsymbol{\xi}^{\boldsymbol{\alpha}}) = F(\boldsymbol{\xi}^{\boldsymbol{\alpha}}) - k_{\mathrm{B}}T \sum_{i} \left(\lambda_i^H N_{\mathrm{H},i}^{\alpha} + \lambda_i^C N_{\mathrm{C},i}^{\alpha}\right) \quad \text{(S11)},$$

where $N_{\mathrm{H},i}^{\alpha}$ and $N_{\mathrm{C},i}^{\alpha}$ are number of amide H-bonds and heavy atoms contacts averaged within bin $\alpha$. Fig. 3CD in the main text was obtained by applying HDXer to the four-dimensional free energy landscape as a function of $\xi_6$, $\xi_{13}$, $\xi_{16}$ and $\xi_{17}$ (see methods in the main text and Fig. S2). To prevent a population bias we rescaled the landscape to have 50% probability for both OF and IF states (defined with 2 kcal/mol cutoff from each minima). We removed intermediate and transition region of the landscape (Fig. 2A) as preliminary reweighting trials show that are not significantly affected by the reweighting. The HDX-MS data for WT and 5L6-8 NCX_Mj used in the HDXer analysis, refer to the measurements reported by Giladi et al.(18), obtained under saturating $Na^+$ concentration (more consonant with the simulation setup). This data consists of deuteration levels at 15 s, for 12 protein fragments (40-46, 47-59, 53-59, 194-204, 203-214, 205-214, 215-230, 215-237, 219-250, 238-245, 238-250, 259-264), which comprise portions of the $\alpha$-repeats (TM2, TM7 and TM8, Fig. 3A).

*Extraction of conformational ensembles for OF and IF states from free-energy basins*

The conformational ensembles considered in comparative analysis of OF, intermediate and IF states were extracted from the BE-META simulations, in reference to the features of the free-energy landscapes. Specifically, the OF state is defined by $\xi_6 \leq 1.3$ and $\xi_{13} \leq 1.55$; the intermediate state by $1.4 < \xi_6 < 1.7$ and $1.35 < \xi_{13} < 1.7$ and $1.4 < \xi_{16} < 1.7$; and the IF state by $\xi_6 > 1.45$ and $\xi_{13} > 1.7$. For the calculation of mean values of a certain descriptor for each ensemble, we included only configurations within 1 kcal/mol from the lowest free-energy configuration in each case. These configurations were selected based on a Metropolis-Monte Carlo acceptance criteria, which accounts for the free energy and thus produces an unbiased sample.

*Construction of alternative ion-occupancy states*

The states with 2 $Na^+$ ions bound to NCX_Mj were generated from the states with 3 $Na^+$ ions bound through a slow alchemical transformation that annihilates the ion bound at either the $S_{ext}$ or the $S_{int}$ site, over a 40-ns MD trajectory. A $Cl^-$ ion in the bulk solution was concurrently annihilated to maintain neutrality. The apo states (i.e., no ions) were constructed from the equilibrated states with 2 $Na^+$ ions bound by gradually annihilating both ions, again over a 40-ns MD trajectory, together with two $Cl^-$ ions in the bulk solution. The states with 2 $H^+$ bound were generated from the states with 3 $Na^+$ bound by gradually annihilating the $Na^+$ ions from the binding sites, and a $Cl^-$ ion from the bulk, while creating protonated forms of E54 and E213, over a 100-ns MD simulation. Once initial configurations had been obtained, trajectories ranging from 1 to 2 μs were generated for all occupancy states before initiating the enhanced-sampling simulations.

### Supporting Tables

**Table S1**. Gaussian widths ($\sigma$), path collective variables smoothing parameters ($\lambda$) and combinations of reaction coordinates used in each replica of the BE-META simulations of the alternating-access transition of NCX_Mj with either 3 $Na^+$ ions or 1 $Ca^{2+}$ ion bound.[a]

| | $\sigma$ | $\lambda\ [nm^{-2}]$ | Unbiased | $\xi_1$ | $\xi_6$ | $\xi_7$ | $\xi_{14}$ | $\xi_{16}$ |
|---|---|---|---|---|---|---|---|---|
| $\xi_1$ | 0.01 | 7.5 | - | - | - | - | - | - |
| $\xi_2$ | 0.01 | 5 | - | 1, 17, 33, 49 | - | - | - | - |
| $\xi_3$ | 0.013 | 7.5 | - | 2, 18, 34, 50 | - | - | - | - |
| $\xi_4$ | 0.027 | 57 | - | 3, 19, 35, 51 | - | - | - | - |
| $\xi_5$ | 0.027 | 80 | - | 4, 20, 36, 52 | - | - | - | - |
| $\xi_6$ | 0.01 | 6 | - | 5, 21, 37, 53 | - | - | - | - |
| $\xi_7$ | 0.01 | 5.6 | - | - | - | | - | - |
| $\xi_8$ | 0.01 | 9.7 | - | - | - | 11, 27, 43, 59 | - | - |
| $\xi_9$ | 0.013 | 5.5 | - | 6, 22, 38, 54 | - | - | - | - |
| $\xi_{10}$ | 0.013 | 8 | - | 7, 23, 39, 55 | - | - | - | - |
| $\xi_{11}$ | 0.027 | 60 | - | 8, 24, 40, 56 | - | - | - | - |
| $\xi_{12}$ | 0.027 | 66 | - | 9, 25, 41, 57 | - | - | - | - |
| $\xi_{13}$ | 0.01 | 6.1 | - | 10, 26, 42, 58 | 15, 31, 32, 47, 63, 64 | - | - | - |
| $\xi_{14}$ | 0.013 | 6 | - | - | - | - | | - |
| $\xi_{15}$ | 0.013 | 10 | - | - | - | - | 12, 28, 44, 60 | - |
| $\xi_{16}$ | 0.01 | 5.3 | - | - | - | - | - | - |
| $\xi_{17}$ | 0.013 | 9.5 | - | - | - | - | - | 13, 29, 45, 61 |
| $\xi_{18}$ | 0.01 | 5.9 | - | 14, 30, 46, 62 | - | - | - | - |
| Unbiased | - | - | 16, 48 | - | - | - | - | - |

[a] Gaussian functions were added to the Metadynamics biasing potential on each variable in 5-ps intervals. In the equilibration stage of $Na^+$-bound NCX_Mj, we used 32 replicas, all started from the same conformation. To avoid a large initial bias, the strength of the history-dependent potential of Metadynamics was gradually raised during the first 500 ns. Specifically, the Gaussians height was set to 0.25 kJ.mol for the first 360 ns and then gradually incremented to 1 kJ/mol from 360 ns to 500 ns. After 1 μs (for each replica) we duplicated the number of replicas and lowered the Gaussians height to 0.5 kJ/mol for the remaining 200 ns of sampling. For $Ca^+$-bound NCX_Mj, the Gaussians height was set to 0.5 kJ/mol.

**Table S2.** Gaussian widths ($\sigma$) and temperature bias (ΔT) used in each of the replicas in the bias-exchange well-tempered Metadynamics simulations of opening and occlusion of NCX_Mj[a].

| Replica | Collective variables biased | $\sigma(\xi_{int})$ [nm] | $\sigma$ ($\xi_{Ca}$) [nm] | $\sigma$ ($\xi_{TM1,2-7}$) | $\sigma$ ($\xi_{TM2-8}$) | 298K+ΔT(K) |
|---|---|---|---|---|---|---|
| 1 | $\xi_{int}$, $\xi_{Ca}$ | 0.025 | 0.025 | - | - | 4470 |
| 2 | $\xi_{int}$, $\xi_{Ca}$ | 0.025 | 0.025 | - | - | 4470 |
| 3 | $\xi_{int}$, $\xi_{Ca}$, $\xi_{TM2-8}$ | 0.117 | 0.117 | - | 0.233 | 2214 |
| 4 | $\xi_{int}$, $\xi_{Ca}$, $\xi_{TM2-8}$ | 0.100 | 0.100 | - | 0.200 | 1831 |
| 5 | $\xi_{int}$, $\xi_{Ca}$, $\xi_{TM2-8}$ | 0.083 | 0.083 | - | 0.166 | 1447 |
| 6 | $\xi_{int}$, $\xi_{Ca}$, $\xi_{TM2-8}$ | 0.067 | 0.067 | - | 0.133 | 1064 |
| 7 | $\xi_{int}$, $\xi_{Ca}$, $\xi_{TM2-8}$ | 0.050 | 0.050 | - | 0.100 | 681 |
| 8 | unbiased | - | - | - | - | - |
| 9 | $\xi_{TM1,2-7}$, $\xi_{TM2-8}$ | - | - | 0.017 | 0.050 | 4470 |
| 10 | $\xi_{TM1,2-7}$, $\xi_{TM2-8}$ | - | - | 0.017 | 0.050 | 4470 |
| 11 | $\xi_{int}$, $\xi_{Ca}$, $\xi_{TM1,2-7}$ | 0.117 | 0.117 | 0.082 | - | 2214 |
| 12 | $\xi_{int}$, $\xi_{Ca}$, $\xi_{TM1,2-7}$ | 0.100 | 0.100 | 0.070 | - | 1831 |
| 13 | $\xi_{int}$, $\xi_{Ca}$, $\xi_{TM1,2-7}$ | 0.083 | 0.083 | 0.058 | - | 1447 |
| 14 | $\xi_{int}$, $\xi_{Ca}$, $\xi_{TM1,2-7}$ | 0.067 | 0.067 | 0.046 | - | 1064 |
| 15 | $\xi_{int}$, $\xi_{Ca}$, $\xi_{TM1,2-7}$ | 0.050 | 0.050 | 0.035 | - | 681 |
| 16 | $\xi_{TM1,2-7}$, $\xi_{TM2-8}$ | - | - | 0.017 | 0.050 | 4470 |

[(a)] Gaussian functions were added to the biasing potential acting on each variable at 5 ps intervals and were initially 2 kJ/mol in height. The height of these Gaussians functions was progressively scaled down as prescribed by the well-tempered Metadynamics method(19). Note the specified 'temperature bias' is not the simulation temperature, which was in all cases 298 K. Following a previous study (2), we used this scheme for an efficient exploration of the multidimensional collective-variable space: replicas with larger values of the temperature-bias and Gaussian widths are able to access high free-energy regions, albeit at low spatial resolution. Replicas with smaller values primarily sample low free-energy regions but at high spatial resolution.
